# Single-cell genotype-phenotype mapping identifies therapeutic vulnerabilities in VEXAS syndrome

**DOI:** 10.1101/2024.05.19.594376

**Authors:** Saravanan Ganesan, Rebecca M. Murray, Jesus Sotelo, Elliot O. Eton, Kouhei Takashima, Theo Botella, Kai Beattie, Alyssa C. Indart, Nada Chraiki, Carolyne Croizier, Franco Izzo, Catherine Potenski, Samuele Marro, Zhijie Wu, Shouguo Gao, Neal S. Young, John D. Blair, Rahul Satija, Benjamin Terrier, Mael Heiblig, Ivan Raimondi, Eirini P Papapetrou, Pierre Sujobert, Olivier Kosmider, David B. Beck, Dan A. Landau

## Abstract

Somatic evolution leads to the emergence of clonal diversity across tissues with broad implications for human health. A striking example of somatic evolution is the VEXAS (Vacuoles E1 enzyme X-linked Autoinflammatory Somatic) syndrome, caused by somatic *UBA1* mutations in hematopoietic stem cells (HSCs), inducing treatment-refractory, systemic inflammation. However, the mechanisms that lead to survival and expansion of mutant HSCs are unknown, limiting the development of effective therapies. The lack of animal or cellular models of *UBA1*-mutant HSCs has hindered such mechanistic understanding, mandating analysis of primary human VEXAS samples, which harbor admixtures of wild-type and *UBA1*-mutant HSCs. To address these challenges, we applied single-cell multi-omics to comprehensively define mutant *UBA1*-induced transcriptome, chromatin accessibility and signaling pathway alterations in VEXAS patients, allowing for the direct comparison of mutant versus wild-type cells within the same environment. We confirmed the expected enrichment of *UBA1*^M41V/T^ mutations in myeloid cells, and additionally discovered that these mutations were also prevalent in Natural Killer (NK) cells in VEXAS patients, providing new insights into disease phenotypes. Through mapping genotypes to molecular phenotypes, including transcriptome, chromatin accessibility, cell surface protein or intracellular protein profiles, in HSCs, we found that *UBA1*^*M41V/T*^-mutant cells showed an increased inflammation signature (interferon alpha and gamma response pathways), as well as activation of unfolded protein response (UPR) via pro-survival, but not pro-apoptotic, mediators of the PERK pathway, compared to *UBA1* wild-type HSCs. Ex vivo validation experiments showed that inhibiting *UBA1* in normal CD34+ or using *UBA1*-mutant HSCs led to PERK pathway up-regulation, increased myeloid differentiation and cell survival, which was reversed by PERK inhibition. Thus, we demonstrated that human VEXAS HSCs show cell-intrinsic inflammatory phenotypes and survive the proteomic stress caused by compromised ubiquitination through PERK-mediated activation of the UPR. Together, these analyses nominate PERK pathway inhibition as a potential new therapeutic strategy for eradicating the VEXAS-inducing clone, demonstrating the power of single-cell multi-omics profiling of primary patient sample to enable genotype-to-phenotype somatic mapping for the discovery of novel candidates for clinical intervention.

## Introduction

Recent discoveries have radically altered our understanding of the human genome, revealing that each human is composed of a multitude of heterogeneous clones, representing not one genome but many. These clonal expansions are fueled by somatic driver mutations that confer a selective advantage and can be observed even in normal tissues^1–5^ as well as in non-malignant disease^6–10^. Non-malignant somatic evolution has been best studied in the hematopoietic system, where it was shown that clonal expansions are nearly ubiquitous in hematopoietic stem cells (HSCs), resulting in highly prevalent clonal hematopoiesis of indeterminate potential (CHIP) by the age of 70^11^, which has been linked to various cardiovascular, metabolic and auto-immune diseases^12^. The frequency of clonal outgrowths, together with their links to disease, raises central biological questions about how clonal expansions impact tissue function across physiological and pathological contexts.

The recently described vacuoles E1 enzyme X-linked autoinflammatory somatic (VEXAS) syndrome represents a novel somatic disorder of the hematopoietic system^13^. This adult-onset disease is caused by hypomorphic, somatic mutations in the ubiquitin-activating enzyme E1 (*UBA1*) gene (p.Met41) originating in HSCs. These somatic mutations lead to accumulation of misfolded proteins, activation of the unfolded protein response (UPR) and ultimately systemic inflammation accompanied by severe clinical symptoms with overall five-year survival of only 63%^14^. *UBA1* (through isoforms *UBA1a* and *UBA1b*) encodes the major E1 enzyme responsible for initiating all cellular ubiquitination, thereby regulating most cellular pathways^15^. *UBA1*-mutant cells instead produce a third isoform (*UBA1c*) that fails to properly ubiquitinate, and thus degrade proteins, leading to inflammation and biasing differentiation of HSCs towards the myeloid lineage^13^. These aberrations are associated with an increased risk of overt hematological neoplasia including myelodysplastic syndrome (MDS) and plasma cell dyscrasia. Current VEXAS therapies aim to curb inflammation rather than target the disease-initiating mutant HSCs^16^, mainly due to our lack of biological understanding of the survival mechanisms of *UBA1*-mutated HSCs. Previous studies focused on inflammation and myeloid bias in hematopoietic stem and progenitor cells (HSPCs)^17^ and differentiated monocytes^18^. However, they lacked sufficient resolution to study the less abundant HSC population to address the pathological effects of *UBA1* mutation. Notably, mouse models are lacking and *UBA1*-mutant cells are challenging to study in vitro due to inability of the mutant stem and progenitor cells to proliferate and expand. Therefore, direct analysis of VEXAS patients is needed in order to characterize the cellular and molecular features of this disease, especially the phenotypes of the disease-initiating HSCs, which remain poorly understood. Key outstanding questions include: (i) how do mutant HSCs survive and clonally expand despite high proteome toxicity; (ii) does inflammation and myeloid bias arise from the HSCs or from other progenitor cells; and (iii) do phenotypic changes driven by somatic *UBA1* mutations represent cell-intrinsic vulnerabilities that could be exploited for the elimination of disease-initiating HSCs?

The study of somatic clonal expansions in humans is limited by a central challenge - mutant and wild-type cells are admixed. To overcome this limitation, single-cell multi-omics techniques^19–23^ allow for high-throughput analysis of both wild-type and mutant cells, in order to characterize the mutant cell-specific changes to the transcriptome, epigenome and proteome. Importantly, this approach turns the admixture of mutant and wild-type cells in somatic tissue from a limitation to an advantage, enabling the direct comparison of mutant and wild-type cells within the same individual, overcoming patient-specific confounders in human studies^24,20^. Crucially, as the single-cell multi-omics approach compares mutant and wild-type cells within the same environment, it is poised to identify mutant-specific differences that can serve as a therapeutic vulnerability to eliminate mutant, but not wild-type cells.

Here, to address outstanding mechanistic questions relating to VEXAS pathology and identify factors that could potentially be targeted for eradication of the disease-initiating HSCs, we apply multi-omic single-cell techniques, including Genotyping of Transcriptome (GoT) plus cellular indexing of transcriptomes and epitopes (CITE-seq)^25^ and Genotyping of Targeted Loci with Chromatin Accessibility (GoT-ChA)^23^ plus cell surface or intracellular (Phospho-Seq)^26^ protein profiling by sequencing directly to VEXAS patient samples. High-resolution mapping of phenotypes across transcriptome, cell surface protein profiling, intracellular protein profiling and chromatin accessibility of *UBA1*^*M41V/T*^ vs. *UBA1* wild-type cells shows mutation enrichment in myeloid cells, as well as in natural killer (NK) cells, which could be functionally impacted by the *UBA1*^*M41V/T*^ mutation. Analysis of HSCs reveals myeloid priming of mutated cells, with increased inflammation signatures and global translational downregulation accompanied by increased ATF4 target gene enrichment, implicating the PKR-like endoplasmic reticulum kinase (PERK) branch of the UPR pathway. Functional validation of the specific role of the PERK pathway on the fitness of UBA1-inhibited HSCs shows that inhibition of the PERK pathway reduces the expression of *ATF4* and correlates with the reduced viability of CD34+ cells. Altogether, we demonstrate that VEXAS HSCs are myeloid biased, show a cell-intrinsic inflammatory phenotype and tolerate proteome toxicity through activation of the anti-apoptotic arm of the PERK pathway as a part of UPR, providing a key survival mechanism to *UBA1*^*M41V/T*^-mutated HSCs. Our results suggest that targeting the PERK pathway in HSCs may represent a new strategy for the treatment of VEXAS syndrome, and underscore the value of single-cell genotype-to-phenotype mapping in human somatic mosaicism.

## Results

### Single-cell analysis in VEXAS syndrome reveals cell type-specific inflammatory and stress responses

To analyze the impact of somatic mutation as a function of cell type in VEXAS and uncover molecular mechanisms of mutation-driven clonal outgrowth in somatic evolution, we combined GoT^19^ – a droplet-based method for joint single-cell (sc)RNA-seq and capture of somatic mutations – with epitope profiling via CITE-seq^25^, and applied this to CD34+ enriched bone marrow progenitor cells (**Supp. Fig. 1a**) from VEXAS patients (n = 10) and healthy controls (n = 4, cohort and sampling details can be found in **Supp. Table 1**). Following sequencing and quality control filtering, we obtained 83,107 cells across the 10 VEXAS samples, as well as 54,368 cells from the 4 healthy donors. To elucidate potential molecular mechanisms underlying VEXAS pathology, we first integrated the data across all VEXAS samples and clustered cells according to transcription information, agnostic to genotype (**Fig. 1a**). We annotated cell types by transferring cell type labels from the Azimuth bone marrow reference and then confirmed cluster identities using surface protein data, identifying the expected clusters reflecting hematopoietic differentiation. We first compared cells from VEXAS patients (irrespective of genotyping) to healthy control cells (**Fig. 1b, Supp. Fig. 2**). In agreement with previous reports^13,17^, we observed that VEXAS HSCs exhibit a myeloid bias, with higher myeloid differentiation scores (Methods) compared to healthy controls (*P* < 2.22 × 10^−16^; **Fig. 1c**). Differential gene expression in VEXAS versus control HSCs (**Fig. 1d**), as well as enriched gene sets across cell type clusters (**Fig. 1e, Supp. Table 3**), revealed enrichment in inflammation, proteome stress and unfolded protein response pathways and depletion for translation pathways compared to controls, consistent with protein degradation pathway disruption mediated by *UBA1* mutation (**Fig. 1f**).

**Figure 1:**
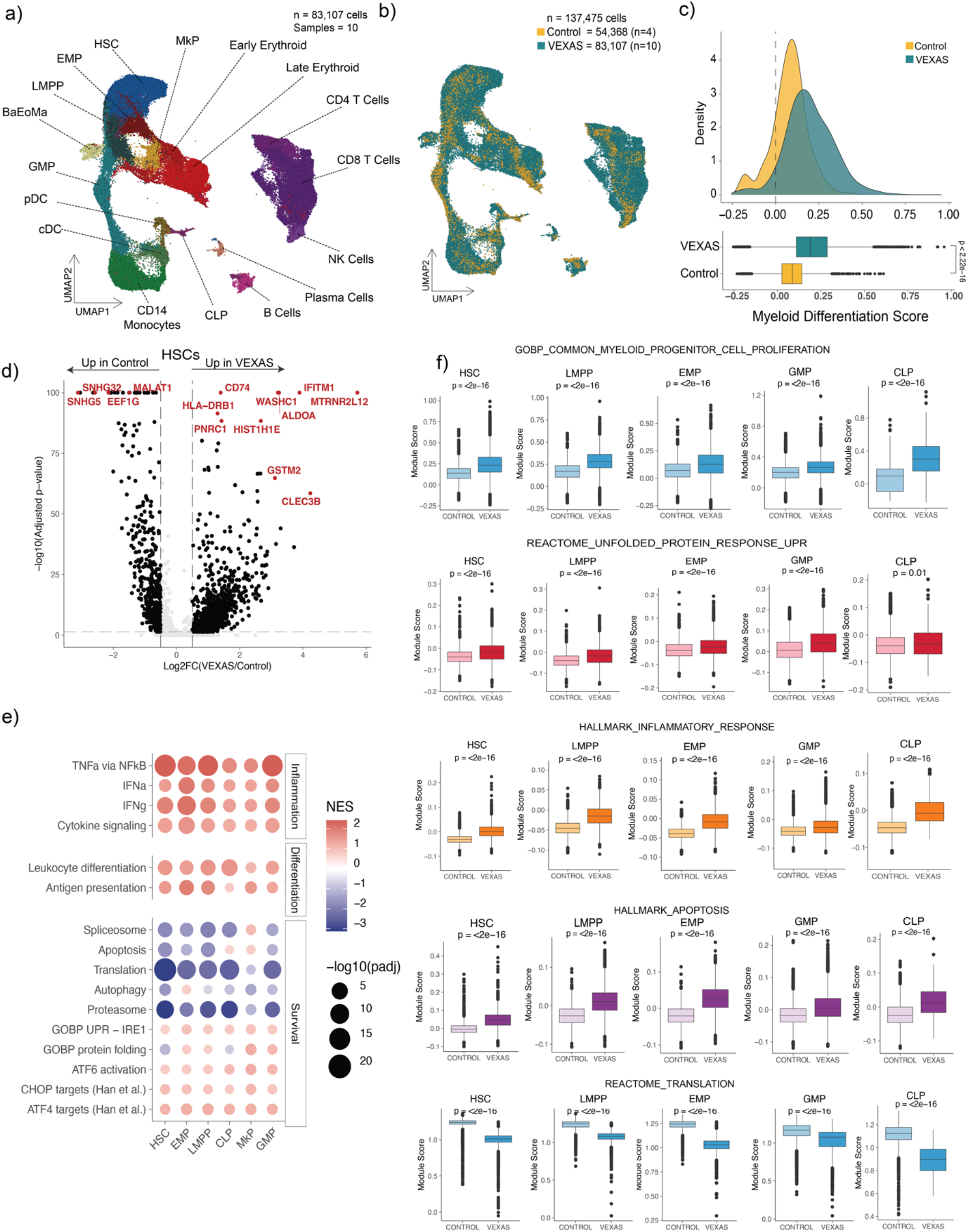
Single-cell analysis of CD34+ compartment in VEXAS syndrome reveals cell type-specific inflammatory and stress responses. **(a)** Gene expression UMAP from patient samples (n = 10 patients, 83,107 cells) with annotated clusters (Methods). HSC, hematopoietic stem cells; EMP, erythro-myeloid progenitors; LMPP, lymphoid-myeloid pluripotent progenitors; MkP, megakaryocytic progenitors; BaEoMa, basophil/eosinophil/mastocyte progenitors; GMP, granulocyte monocyte progenitors; CLP, common lymphoid progenitors; pDC, plasmacytoid dendritic cells; cDC, classical dendritic cells; NK, natural killer cells. **(b)** UMAP from (**a**) with cells from VEXAS samples (green, n = 10 patients; n = 82,107 cells) overlaid with cells from healthy controls (yellow, n = 4 individuals; n = 54,368 cells). **(c)** Myeloid differentiation scores of HSC population from (**b**) (green, VEXAS patients; yellow, healthy controls, *P* < 2.22 × 10^−16^, two-sided Wilcoxon rank sum test). Myeloid differentiation scores were calculated by scoring each cell for enrichment of a myeloid differentiation gene module (Supp. Table 3, Methods). Error bars represent the range, boxes represent the interquartile range and lines represent the median. **(d)** Volcano plot illustrating differential gene expression scores between control (n = 3,976 cells) and VEXAS (n = 12,739 cells) HSCs. Horizontal dotted line represents *P* = 0.05; Bonferroni-adjusted; vertical dotted lines represent absolute log2FC > 0.5. Genes that are upregulated in VEXAS and control samples are highlighted. Random downsampling was performed to 200 HSCs per donor to minimize patient-specific effects. **(e)** Dot plot showing gene set enrichment scores across cell type clusters. Color scale indicates the mean normalized enrichment score (NES) difference between VEXAS and control cells, and dot size indicates Benjamini-Hochberg-adjusted p-value. **(f)** Gene set module scores in VEXAS vs control HSC, LMPP, EMP, CLP and GMPs cells for myeloid, protein stress response, inflammation, apoptosis and translation pathways. Cell numbers are found in Supp. Table 4. Error bars represent the range, boxes represent the interquartile range and lines represent the median. Two-sided Wilcoxon rank sum test.

### *UBA1* mutation impacts the function of differentiated cell types in VEXAS

In our GoT approach we amplify the locus of interest (**Supp. Fig. 1b, c)**, and integrate genotyping with scRNA-seq data as well as CITE-seq^25^ antibody-derived tag (ADT) information via shared cell barcodes (**Fig. 2a, Supp. Fig. 3**), allowing for the simultaneous profiling of genotype, transcriptome and cell-surface protein expression (Methods, **Fig. 2b, Supp. Fig. 4**). We genotyped *UBA1* in 16,676 out of 83,107 total cells (median 17.16%; range 7.10-32.49% across donors, **Supp. Table 4**), achieving improved genotyping efficiency compared with a non-targeted approach^17^. Compared to total bone marrow mononuclear cells (BMMNCs), the HSPCs (CD34+ cells) showed a higher genotyping efficiency (n = 13,121 cells genotyped out of 38,591 CD34+ cells, median – 31.92 %, range 13.47% - 60.58%; **Supp. Table 5**). Using the same samples reported by Wu et al.^17^, we improved the median genotyping efficiency of *UBA1* from 3.98% (range 2.5 - 5.76%) to 18.08% (range 16.47-22.94%; **Supp. Fig. 4d, Supp. Table 5**) for BMMNCs and from 12.39% (range 2.99% - 24.82%) to 44.58% (range 29.48% - 60.58%; **Supp. Table 5**) for HSPCs, allowing us to capture more genotyped HSCs (from 199 to 1,603 cells out of 5,361 HSCs captured). In line with previous reports^13,17^, we confirmed that *UBA1* mutations are mostly restricted to the myeloid lineage and are depleted in lymphoid cells (**Fig. 2b,c**), consistent with our data showing a myeloid bias of VEXAS samples compared with normal controls. Pseudo-temporal ordering of cells along differentiation trajectories showed a substantial decrease in the fraction of mutant cells along lymphoid differentiation, while mutant cell fraction remained high across myeloid differentiation (**Fig. 2d, Supp. Fig. 4e**).

**Figure 2:**
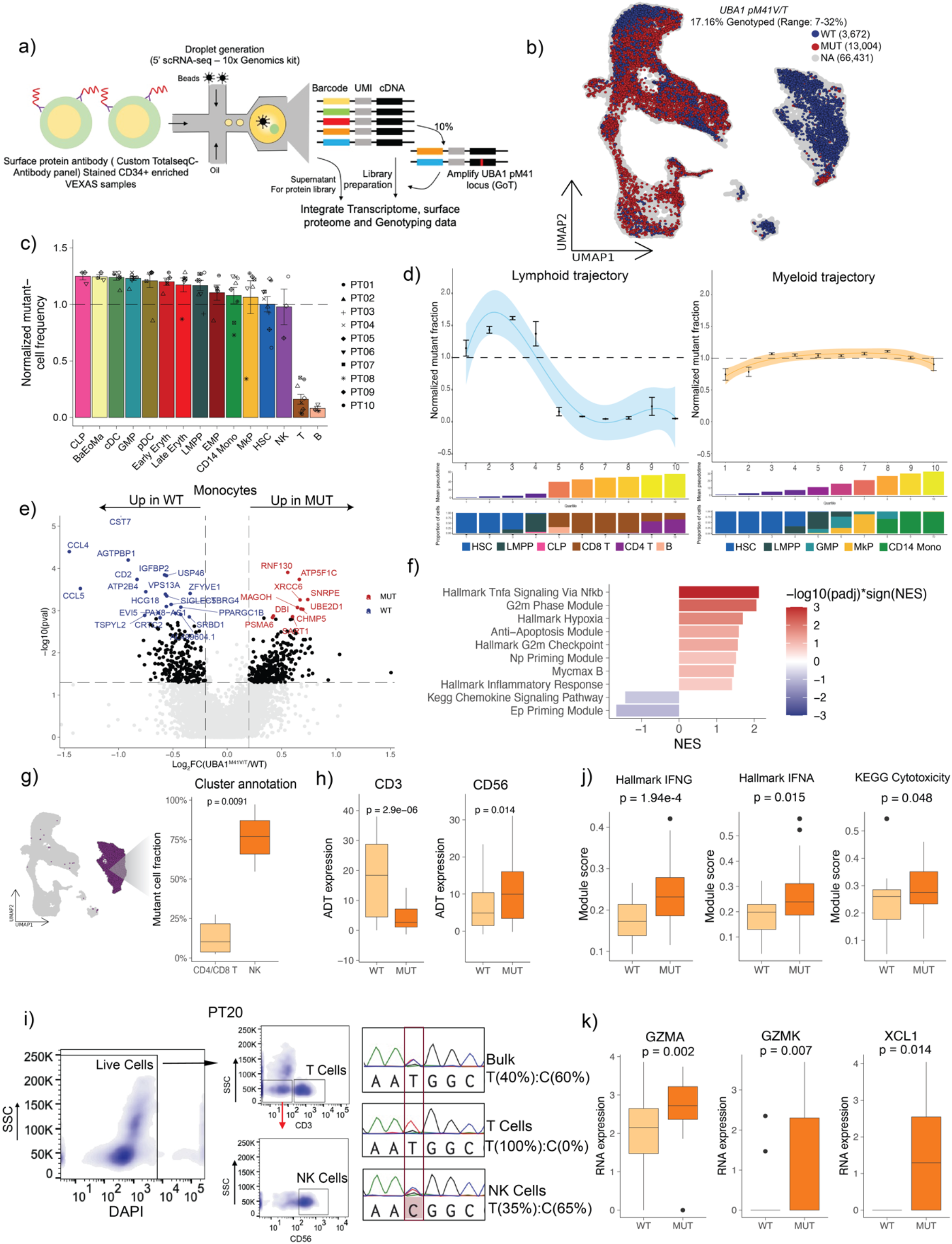
Transcriptomic landscape of *UBA1*-mutant differentiated cell types. **(a)** Schematic of CITE-seq + Genotyping of Transcriptomes (GoT). **(b)** GoT genotyping of *UBA1* projected on the gene expression UMAP from Fig. 1a. Wild-type (WT, blue; n = 3,672 cells), *UBA1*-mutant (MUT, red; n = 13,004 cells) and not assignable (NA, grey; n = 66,431 cells) are shown. **(c)** Mutant cell fraction per cell cluster across patients (n = 10 patients; see Supp. Table 4 for cluster cell numbers per patient). Error bars represent standard error of the mean. **(d)** Normalized mutant fraction along lymphocytic (left) and myeloid (right) pseudotime across VEXAS patient samples (n = 10 patients). Pseudotime was divided in 10 quantiles; each point represents the mean fraction of mutant cells, error bars indicate standard error across samples, lines indicate the fit and shadowed areas represent the 95% confidence interval of the generalized additive model (upper panels). **(e)** Volcano plot illustrating differential gene expression scores between wild-type (n = 52 cells) and mutant (n = 373 cells) CD14+ monocytes in VEXAS patients (n = 5). Horizontal dotted line represents *P* = 0.05; vertical dotted lines represent absolute log2FC > 0.2. Genes that are upregulated in wild-type cells (blue) and mutant cells (red) are highlighted. Linear mixture model (LMM) modeling patient sample as a random effect to account for inter-patient variability followed by likelihood ratio test (Methods). **(f)** Gene set enrichment analysis illustrating enrichment of inflammation (TNFα signaling via NF-κB and Inflammatory response) pathways in *UBA1*-mutant CD14+ monocytes compared to wild-type CD14+ monocytes. Preranked gene set enrichment analysis was performed via the fgsea package (Methods). **(g)** Left, UMAP showing cells in the lymphoid cluster (purple, n = 27,225 cells). Right, mutant cell fraction of genotyped cells in the CD4/CD8 T (n = 2,403 cells) and NK (n = 91 cells) cell clusters. Mutant cell fraction is aggregated by patient, and only patients with >10 genotyped cells were included (n = 3 for NK cells, n = 9 for T cells). Error bars represent the range, boxes represent the interquartile range and lines represent the median. Two-sided Wilcoxon rank sum test. **(h)** Analysis of T cell and NK cell surface marker expression levels in mutant (MUT, n = 32) versus wild-type (WT, n = 79) cells with surface protein expression in the lymphoid cluster. CD3 is a marker of CD4/CD8 T cells and CD56 is a marker of NK cells. Error bars represent the range, boxes represent the interquartile range and lines represent the median. Two-sided Wilcoxon rank sum test. **(i)** Sanger sequencing traces of *UBA1* from a bulk, flow-sorted T cells (CD3) or flow-sorted NK cells (CD56) from a VEXAS patient sample. Sorted NK cells have the UBA1.pM41T(c.122 T>C) mutation. **(j)** Gene set module scores in *UBA1*-mutant vs wild-type NK cells (n = 22 wild-type; n = 64 mutant cells) for Interferon-gamma, Interferon-alpha and Cytotoxicity pathways. Boxes represent the median, bottom and top quartiles, and whiskers correspond to 1.5 the interquartile range. Two-sided Wilcoxon rank sum test. **(k)** Gene expression levels in *UBA1*-mutant vs wild-type NK cells (n = 22 wild-type; n = 64 mutant cells) for *GZMA, GZMK* and *XCL1*. Boxes represent the median, bottom and top quartiles, and whiskers correspond to 1.5 × the interquartile range. Two-sided Wilcoxon rank sum test.

First, to benchmark the analysis, we performed differential gene expression analysis between *UBA1*-mutant and wild-type monocytes, which revealed upregulation of genes involved in inflammation and misfolded protein stress in *UBA1*-mutant cells (**Fig. 2e, Supp. Table 6**), and gene set enrichment analysis highlighted tumor necrosis factor (TNF) signaling via NF-κB, anti-apoptosis genes and inflammatory response pathways in *UBA1*-mutant monocytes (**Fig. 2f, Supp. Table 7**), as expected^18^. Despite the depletion of *UBA1*-mutated lymphoid cells, we observed a previously unreported high frequency of mutant cells in the cluster annotated as NK cells (**Fig. 2b,c**). Indeed, while T cells were mostly wild type (mean mutant cell fraction 0.103 ± 0.102 standard error), NK cells were largely mutant (median mutant cell fraction 0.769 ± 0.212 standard deviation; *P* = 9.1 × 10^−3^; **Fig. 2g**). We further leveraged the cell-surface protein data to compare the expression levels of CD3+ (marking T cells) and CD56+ (marking NK cells) in wild-type versus *UBA1*-mutant cells from the lymphocyte cluster (highlighted in purple in **Fig. 2g**), observing increased CD56 expression in mutant cells (*P* = 0.014) (**Fig. 2h**). We validated this finding in additional patient peripheral blood samples (n = 6) sorted for T cells and NK cells, where we confirmed presence of the *UBA1* mutation [*UBA1*^*M41T*^(c.122 T>C)] specifically in NK cells (4 out of 6 samples) via Sanger sequencing (see **Fig. 2i** for PT20 results, and **Supp. Fig. 5** for results from PT21-PT25). To explore the possible disease-related consequences of *UBA1* mutation enrichment in NK cells in VEXAS, we analyzed differential gene expression between *UBA1*-mutant and wild-type NK cells. Mutant NK cells had enriched module scores for interferon and cytotoxicity pathways (**Fig. 2j**), suggesting an inflammatory phenotype with enhanced killing activity in mutant NK cells. Consistently, we observed upregulation of granzyme (*GZMA, GZMK*) and chemokine (*XCL1*) genes (**Fig. 2k, Supp. Table 6**), further supporting increased cytotoxic signaling and function in mutant NK cells. Together, these results recapitulate the previously observed myeloid bias of the *UBA1*^*M41V/T*^ mutation in VEXAS patients^13,17^, but additionally uncover the presence of the mutation in NK cells, which show increased inflammation and enhancement in cytotoxic gene signatures compared to wild-type NK cells, potentially impacting VEXAS disease pathology.

### *UBA1* mutation leads to increased inflammation and proteomic stress response in hematopoietic progenitors

We next focused our analyses on early HSCs (n = 3,211 cells genotyped out of 12,739 HSCs, median-24.06%, range 6.51-45.45%; including samples from Wu et al.^17^ for which GoT was performed; **Fig. 3a**), as these are the disease-initiating cells in VEXAS, and previous single-cell work^17^ has been limited in its ability to perform genotype-to-phenotype associations of this relatively underrepresented cell population (199 HSCs genotyped; 43 WT and 156 MUT cells; **Supp. Table 5**). Integration of single-cell *UBA1* genotyping with cell surface protein expression revealed that the mutant HSCs showed enrichment of myeloid markers (CD13, CD33, HLA-DR) and depletion of lymphoid markers (CD49b) compared to wild-type cells (**Fig. 3b,c, Supp. Table 8**), demonstrating that *UBA1* mutation already promotes myeloid bias within early progenitors. Differential gene expression analysis in *UBA1*-mutant versus wild-type HSCs further supported early biased myeloid differentiation (e.g., *MYC, CAT, CD38*)^27–29^ and also revealed increased HSC inflammation (e.g., *IFTM1, IRF1, IL1B*)^30–32^ (**Fig. 3d, Supp. Fig. 6a)**, consistent across patients (**Supp. Fig. 6b,c**). These data comparing mutant and wild-type HSCs from the same microenvironment show that the myeloid bias and inflammation seen in VEXAS are cell-intrinsic phenotypes linked to the mutant allele rather than the inflammatory milieu.

**Figure 3:**
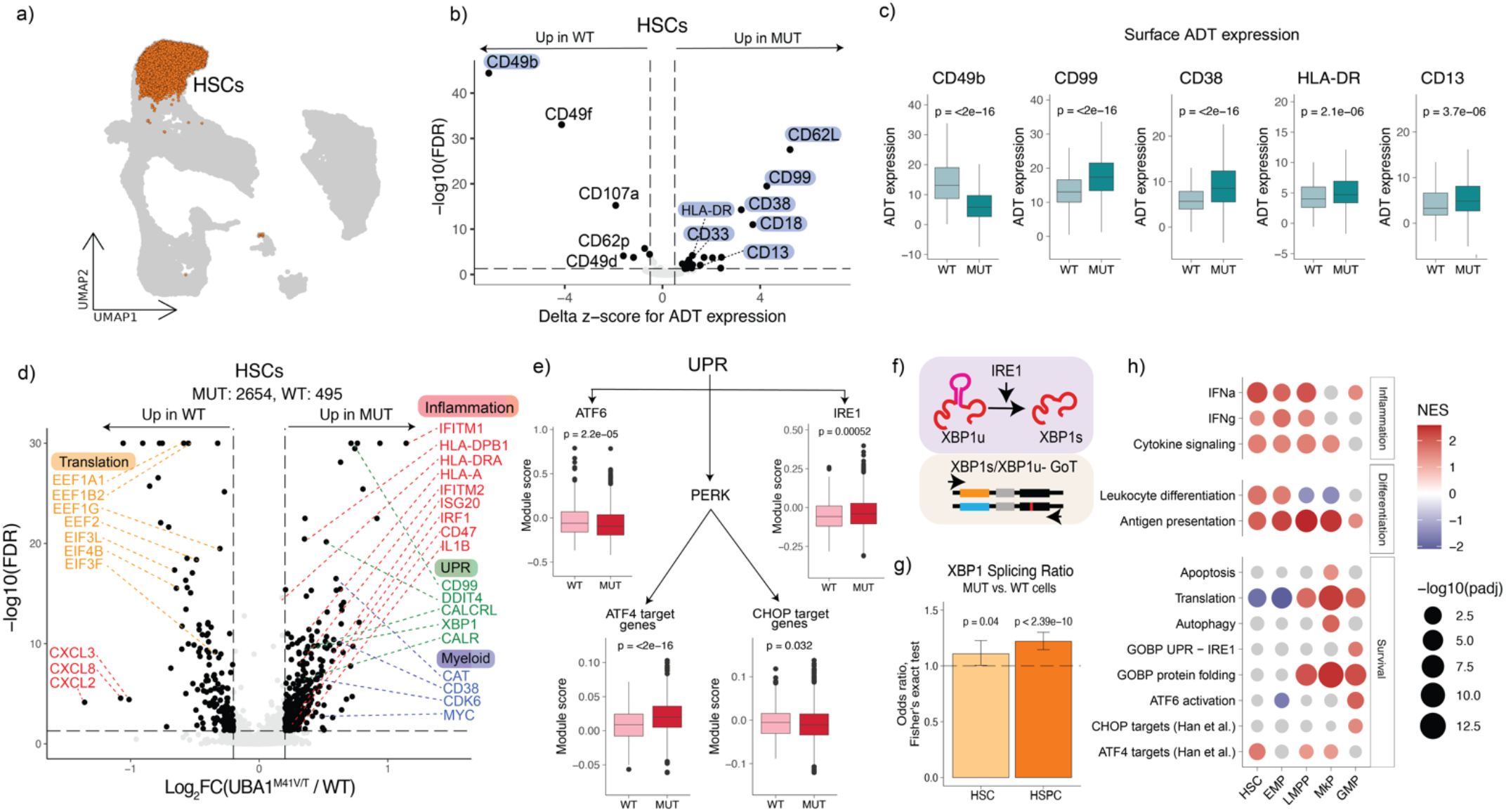
Transcriptomic landscape of *UBA1*-mutant hematopoietic progenitors. **(a)** UMAP from Fig. 2a with HSC cluster highlighted (12,739 HSCs). **(b)** Volcano plot illustrating differential cell surface protein expression [antibody-derived tag (ADT)] scores between wild-type (n = 222 cells) and mutant (n = 1,275 cells) HSCs in VEXAS patients (n = 3). Horizontal dotted line represents FDR = 0.05; vertical dotted lines represent delta z-score > 0.5. Myeloid differentiation genes (blue) are highlighted. Linear mixture model (LMM) modeling patient sample as a random effect to account for inter-patient variability followed by likelihood ratio test (Methods) and Benjamini-Hochberg correction. **(c)** Cell surface protein expression levels in *UBA1*-mutant vs wild-type HSCs for CD49b, CD99, CD38, HLA.DR, and CD13 (n = 222 wild-type cells; n = 1,275 mutant cells). Error bars represent the minimum and maximum values as determined by 1.5 × IQR (outliers not shown), boxes represent the interquartile range and lines represent the median. Two-sided Wilcoxon rank sum test. **(d)** Volcano plot illustrating differential gene expression between wild-type (n = 495 cells) and mutant (n = 2,654 cells) HSCs in VEXAS patients (n = 7). Horizontal dotted line represents FDR = 0.05; vertical dotted lines represent absolute log2FC > 0.2. Genes involved in translation (orange), inflammation (red), unfolded protein response (UPR, green) and myeloid (blue) pathways are highlighted. Linear mixture model (LMM) modeling patient sample as a random effect to account for inter-patient variability followed by likelihood ratio test (Methods) and Benjamini-Hochberg correction. **(e)** Comparison of module scores in *UBA1*-mutant vs wild-type HSCs for GOBP ATF6 and IRE1 pathways, as well as ATF4 target genes and CHOP target genes (n = 503 wild-type; n = 2,708 mutant cells). Error bars represent the range, boxes represent the interquartile range and lines represent the median. Two-sided Wilcoxon rank sum test. **(f)** Schematic and design of GoT targeting for *XBP1* splicing induced by IRE1, with the spliced transcript on right, and the unspliced transcript on the left. The region that is removed during splicing is shown in pink. **(g)** Comparison of spliced (XBP1s) to unspliced (XBP1u) *XBP1* detected in *UBA1*-mutant (n = 2,708 cells) versus wild-type (n = 503 cells) HSCs (yellow bar, left), and *UBA1*-mutant (n = 6,051 cells) versus wild-type (n = 772 cells) HSPCs (orange bar, right) pseudobulked by genotype. P-value determined by Fisher’s exact test. Bars represent the odds ratio, and error bars represent the 95% confidence interval. Dotted line indicates odds ratio of 1. **(h)** Dot plot showing gene set enrichment scores across cell type clusters. Color scale indicates the mean normalized enrichment score (NES) between *UBA1*-mutated and wild-type cells, and dot size indicates Benjamini-Hochberg-adjusted p-value. Modules with padj < 0.1 are greyed out.

Notably, mutated HSCs displayed increased expression of genes involved in the UPR, including *XBP1* and *CALR*, as well as global downregulation of translation compared to wild-type HSCs (**Fig. 3d, Supp. Table 6**), indicating the presence of proteomic stress likely derived from impaired ubiquitination in *UBA1*-mutated HSCs. The UPR is a versatile pathway comprising multiple signaling cascades that are capable of initiating diverse, and even opposing, cellular outcomes, including downregulation of translation, upregulation of autophagy or induction of apoptosis^33,34^. Activation of different arms of the UPR pathway depends on the level and nature of the proteomic stress, as well as the overall state of the cell^35^. Previous reports have shown enrichment for UPR pathway genes as well as impaired ubiquitination in VEXAS patient cells, including HSPCs^13,17^. Given this enrichment of UPR-related genes and pathways, we hypothesized that a potential mechanism for the paradoxical survival and expansion of the ubiquitination-deficient *UBA1*-mutant HSCs could be the induction of a sufficiently cytoprotective UPR that would allow the cells to avoid apoptosis despite proteomic stress.

To better characterize the mechanisms of UPR activation in *UBA1*-mutant HSCs, we analyzed gene sets related to proteomic stress response in mutant versus wild-type cells. We first examined pathways involving the three transmembrane transducers of the UPR: *ATF6, IRE1* and *PERK*^*35*^. We observed no increase in expression of targets of *ATF6*, encoding a transcription factor that is induced by endoplasmic reticulum (ER) stress and activates chaperone genes^36^ (**Fig. 3e**), suggesting that the ATF6 arm of the UPR is not induced in *UBA1*-mutant cells. In contrast, mutant cells showed enrichment for the *IRE1* branch of UPR, which catalyzes the splicing of *XBP1*^*37*^ to promote cell survival^37,38^, indicating that IRE1 activity may be contributing to the mutant HSC phenotype. We sought to directly measure the spliced (XBP1s) and unspliced (XBP1u) *XBP1* transcripts using GoT^19^ in mutant versus wild-type HSCs isolated from VEXAS patients (n = 6, **Fig. 3f**). We designed probes that capture both isoforms of *XBP1* and analyzed the ratio of spliced to unspliced transcript in mutant versus wild-type cells. *UBA1*-mutant HSCs and HSPCs showed higher *XBP1* splicing than wild-type HSCs and HSPCs (*P* = 0.04 and *P* = 2.39x 10^−10^, respectively; **Fig. 3g**, see **Supp. Fig. 6d** for sample-aware permutation analysis, see **Supp. Fig. 6e** for per patient analysis, **Supp. Table 9**), providing further evidence that IRE1 is activated in *UBA1*-mutant cells.

Finally, *PERK* activation can induce apoptosis during sustained unfolded protein stress depending on the context^39,40^, and has previously been shown to be the UPR-mediating pathway in normal HSPCs, contributing to the maintenance of a high-quality HSC pool through the removal of stressed cells^40^. Interestingly, *PERK* has also been shown to activate adaptive rather than cell death pathways in cancer^41^, as well as promote myogenesis during regeneration of skeletal muscles^42^, providing further support that cell state and environment influence UPR outcomes. Consistent with *PERK* activation, gene module scores for the translation pathway were decreased in *UBA1*-mutant HSCs in comparison to wild-type HSCs, indicating a global reduction in protein production (**Supp. Fig. 6a**). We then analyzed downstream signaling of the *PERK* pathway in *UBA1*-mutant versus wild-type HSCs (**Fig. 3e**), observing increased expression of genes targets of ATF4, a stress-induced transcription factor that preferentially promotes cell recovery rather than death upon PERK activation^34^, without any increase in genes targeted by the pro-apoptotic CHOP pathway^43^. These results implicate *IRE1* and *PERK* signaling in promoting the survival, rather than elimination, of *UBA1*-mutant HSCs.

To gain further insights into the pathways enriched in mutant HSPCs across other progenitor clusters, we compared normalized enrichment scores (NES) for myeloid, inflammation, and survival pathways in HSCs to those in erythro-myeloid progenitors (EMPs), lymphoid-myeloid pluripotent progenitors (LMPPs), megakaryocytic progenitors (MkPs) and granulocyte monocyte progenitors (GMPs) (**Fig. 3h**). Notably, mutant MkPs showed no activation of inflammatory programs *(IFNa, IFNg*), but increased expression of translation, autophagy, apoptosis and protein folding gene sets compared to wild-type MkPs. On the other hand, mutant GMP clusters revealed an upregulation of only *IFNa* pathway genes along with increased activation of all three branches of UPR pathway gene sets *IRE1, ATF6* and *PERK* pathways (CHOP targets). Interestingly, this cluster showed only an enrichment for *CHOP* pathway gene sets and no enrichment for *ATF4* gene sets, suggesting that the *PERK* pathway was activated towards a more pro-apoptotic response. Thus, the UPR induced by this mutation in the GMP cluster activates both pro- and anti-apoptotic pathways. Compared to all progenitor clusters, ATF4 target gene set showed the highest upregulation specifically in HSCs, further supporting the involvement of *PERK* activation in contributing to the survival of *UBA1*-mutated HSCs.

### Chromatin accessibility landscape of CD34+ *UBA1*-mutant cells

To determine the regulatory networks that govern the transcriptional impact of the *UBA1* mutation in VEXAS, we integrated our Genotyping of Targeted loci with Chromatin Accessibility profiling (GoT-ChA)^23^ framework with ASAP-seq^44^ to capture genotyping together with chromatin accessibility and cell surface protein profiling (**Fig. 4a**).

**Figure 4:**
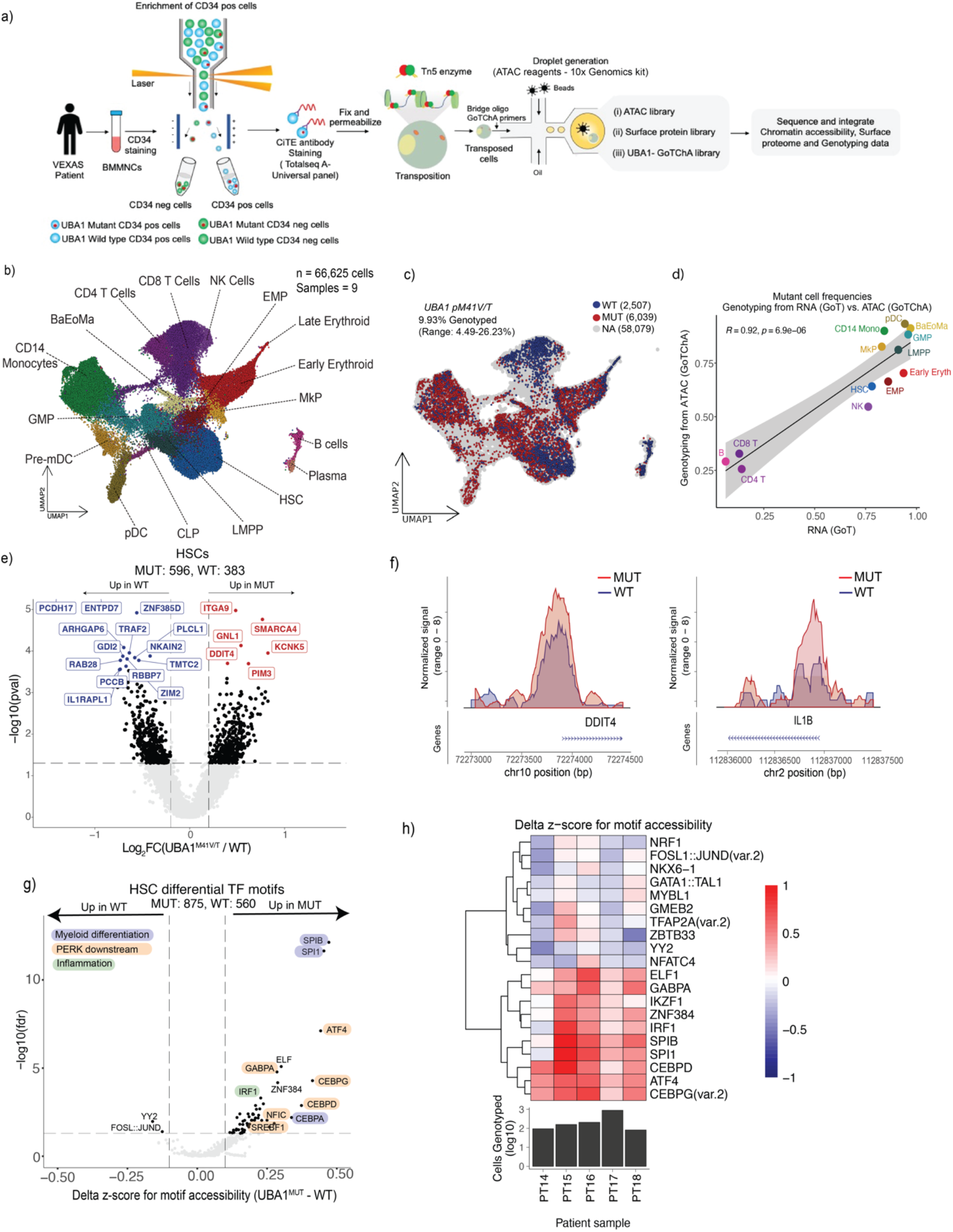
Chromatin accessibility landscape of CD34+ *UBA1*-mutant cells. **(a)** Schematic for GoT-ChA + ASAP-seq approach for simultaneous profiling of genotype, chromatin accessibility and cell surface proteins. **(b)** Chromatin accessibility UMAP showing reconstruction of VEXAS hematopoiesis from human VEXAS samples (n = 66,625 cells) with annotated clusters (Methods). **(c)** GoT-ChA genotyping of *UBA1* projected on the scATAC-seq UMAP from (**c**). Wild-type (WT, blue; n = 2,507 cells), *UBA1*-mutant (MUT, red; n = 6,039 cells) and not assignable (NA, grey; n = 58,079 cells) are shown. **(d)** Pearson correlation between mutant cell fractions for each cluster as determined by either GoT (mRNA-based genotyping) or GoT-ChA (gDNA-based genotyping). Cell types are colored according to the same scheme as in (**b**). Shaded area represents 95% confidence interval. **(e)** Volcano plot illustrating differential gene accessibility scores between wild-type (n = 383 cells) and mutant (n = 596 cells) cells within HSC clusters of VEXAS patients (n = 5). Horizontal dotted line represents p-value = 0.05; vertical dotted lines represent absolute log2FC > 0.2. Top 20 differentially accessible genes when ranked by p-value are highlighted, with genes more accessible in MUT cells in red and in WT cells in blue. Linear mixture model (LMM) modeling patient sample as a random effect to account for inter-patient variability followed by likelihood ratio test (Methods). **(f)** Example normalized accessibility tracks for genes with increased accessibility (*DDIT4*, left; *IL1B*, right) in *UBA1*-mutant (red) versus wild-type (blue) HSCs. **(g)** Volcano plot of the differentially accessible transcription factor motifs (FDR < 0.05 and absolute Δz-score > 0.1; LMM followed by likelihood ratio test and Benjamini-Hochberg correction) between wild-type (n = 560 cells) and *UBA1*-mutant (n = 875 cells) HSCs from VEXAS patients (n = 5). Transcription factors involved in biological pathways including myeloid differentiation (blue), PERK targets (orange) and inflammation (green) are highlighted. The horizontal dotted line represents FDR = 0.05, the vertical dotted lines represent absolute Δz-score > 0.1. **(h)** Top, heatmap of Δz-score for transcription factor motif accessibility in HSCs across five patients. Color scale indicates the mean z-score difference between *UBA1*-mutated and wild-type HSCs. Bottom, genotyped HSC count per patient.

We applied GoT-ChA to primary VEXAS samples (CD34+ cell enriched) from 9 donors and obtained 66,625 cells after quality control filtering (**Fig. 4b, Supp. Fig. 7-9**). We first performed cell clustering based solely on chromatin accessibility data, agnostic to genotype. Doublets were removed and samples were integrated using reciprocal latent semantic indexing (LSI) projection to mitigate batch effects^45^. We identified the expected progenitor subtypes, using surface protein expression as well as cell type labels transferred from Azimuth’s scRNA-seq bone marrow reference via bridge integration^46,47^ (**Supp. Fig. 9**). We genotyped *UBA1* in 8,546 out of 66,625 cells (median 9.93%; range 4.49-26.23% across donors, **Supp. Table 10**), and projected the genotypes onto the differentiation map (**Fig. 4c, Supp. Fig. 10**). To validate that our methods provided comparable levels of genotyping across cell types, we compared the mutant cell fractions of our samples based on GoT RNA-based genotyping vs GoT-ChA gDNA-based genotyping for analyzed cell clusters (**Fig. 4d**). We saw strong correlation between the RNA- and ATAC-based data (R = 0.92, *P* = 6.9 × 10^−6^), confirming the myeloid and NK enrichment and lymphoid depletion of mutant cells.

We next analyzed genes with differential chromatin accessibility between mutant and wild-type cells. Gene accessibility scores were generated using the ArchR software^48^, and genes were filtered for minimum expression levels in both the gene accessibility score data and the RNA expression data of the corresponding cell type (Methods). Focusing on HSCs (1,459 cells genotyped out of 10,303 HSCs; median genotyping efficiency 8%; range 4.46-22.64%), we identified enrichment for genes involved in inflammation and myeloid differentiation in *UBA1*-mutant cells (**Fig. 4e**), consistent with our transcriptome data (**Fig. 3d, Supp. Fig. 6a, Supp. Table 11**). Inspection of peaks at *DDIT4* and *IL1B* regions showed increased accessibility in mutants compared to wild-type HSCs (**Fig. 4f)**, reflecting UPR^49,50^ and inflammation signaling^51^ activation, respectively. Similar to the transcriptome data, gene set enrichment analysis of chromatin accessible regions showed an enrichment for inflammatory pathway genes in mutant cells, compared to wild-type cells (**Supp. Fig. 11, Supp. Table 12**), supporting the model that cell-intrinsic inflammatory signaling in mutant HSCs contributes to disease phenotypes.

We next leveraged the chromatin accessibility profiles to infer transcription factor (TF) activity using DNA binding motif accessibility^52^. Comparing wild-type and *UBA1*-mutated HSCs, we identified TFs with increased motif accessibility (false discovery rate [FDR] < 0.05, Δz-score > 0.1) in mutant cells (**Supp. Table 13**). First, we observed increased motif accessibility of myeloid differentiation factors, including, SPI1, SPIB and CEBPA, consistent with increased myeloid priming. In addition, TFs involved in inflammation (e.g., *IRF1*) had increased motif accessibility in mutant cells, consistent with cell-intrinsic *UBA1* mutation-related inflammatory phenotype. Notably, TFs downstream of PERK activation also had increased motif accessibility in mutant HSCs, including ATF4 (**Fig. 4g**). Differential TF binding motif accessibility was largely consistent across patients, with ATF4 binding motifs showing increased accessibility in all samples with sufficient number of genotyped HSCs (**Fig. 4h**). Together, these results demonstrate that VEXAS *UBA1*-mutant HSCs are epigenetically reprogrammed towards myeloid differentiation, inflammation and UPR activation via the PERK pathway, suggesting that these are cell-intrinsic alterations driven specifically by *UBA1* mutation, providing further support for the development of HSC-targeting strategies for VEXAS therapy.

### PERK-ATF4 signaling is activated in CD34+ *UBA1*-mutant cells

Next, we sought to additionally measure single-cell intracellular protein levels together with genotype to directly validate PERK signaling in mutant cells. To this end, we integrated intracellular protein capture^26^ into our GoT-ChA framework (**Fig. 5a**). We designed an antibody-labeled intracellular protein panel (**Supp. Table 14**) comprising relevant signaling and regulatory factors involved in cell stress response, UPR (e.g., ATF4, CHOP, DAD1), inflammation (e.g., IL-1B, p65, STAT3), myeloid differentiation (e.g., CEBPA, SPI-B, RARA) and cell survival (e.g., MCL1, XIAP). We first standardized this assay using a K562 cell line model treated with UBA1 inhibitor and confirmed assay specificity using a cell line mixing experiment (**Supp. Fig. 12a-c, Supp. Table 15**).

We then applied this assay to two VEXAS patient samples (PT16, n = 12,966 cells; PT17, n = 12,745 cells) and compared protein levels in wild-type vs. *UBA1*-mutant HSPCs (**Fig. 5b**). Even with the addition of the intracellular protein modality into the GoT-ChA framework, we were still able to attain adequate genotyping efficiency (PT16 = 14.8%, PT17 = 26.2%). We assessed intracellular protein levels in both samples and ranked proteins by differential expression between *UBA1*-mutant and wild-type HSCs, (Methods; **Fig. 5b, Supp. Table 15**). In both samples, consistent with our transcriptome and TF motif accessibility data, *UBA1*-mutant cells showed increased intracellular ATF4 protein levels compared to wild-type cells (**Fig. 5b,c**). Conversely, CHOP protein levels were increased in wild-type cells compared to mutant HSCs, indicating that the PERK-mediated UPR is biased away from the pro-apoptotic CHOP pathway in *UBA1*-mutant HSCs. Comparison of these protein levels across other progenitors, including LMPP, EMP, and MkP, showed that ATF4 was consistently upregulated and CHOP was downregulated only in HSCs in both patients tested (**Supp. Fig. 12d**). These results provide further support that PERK-ATF4, but not CHOP, is activated in response to proteomic stress in mutant HSCs in VEXAS.

**Figure 5:**
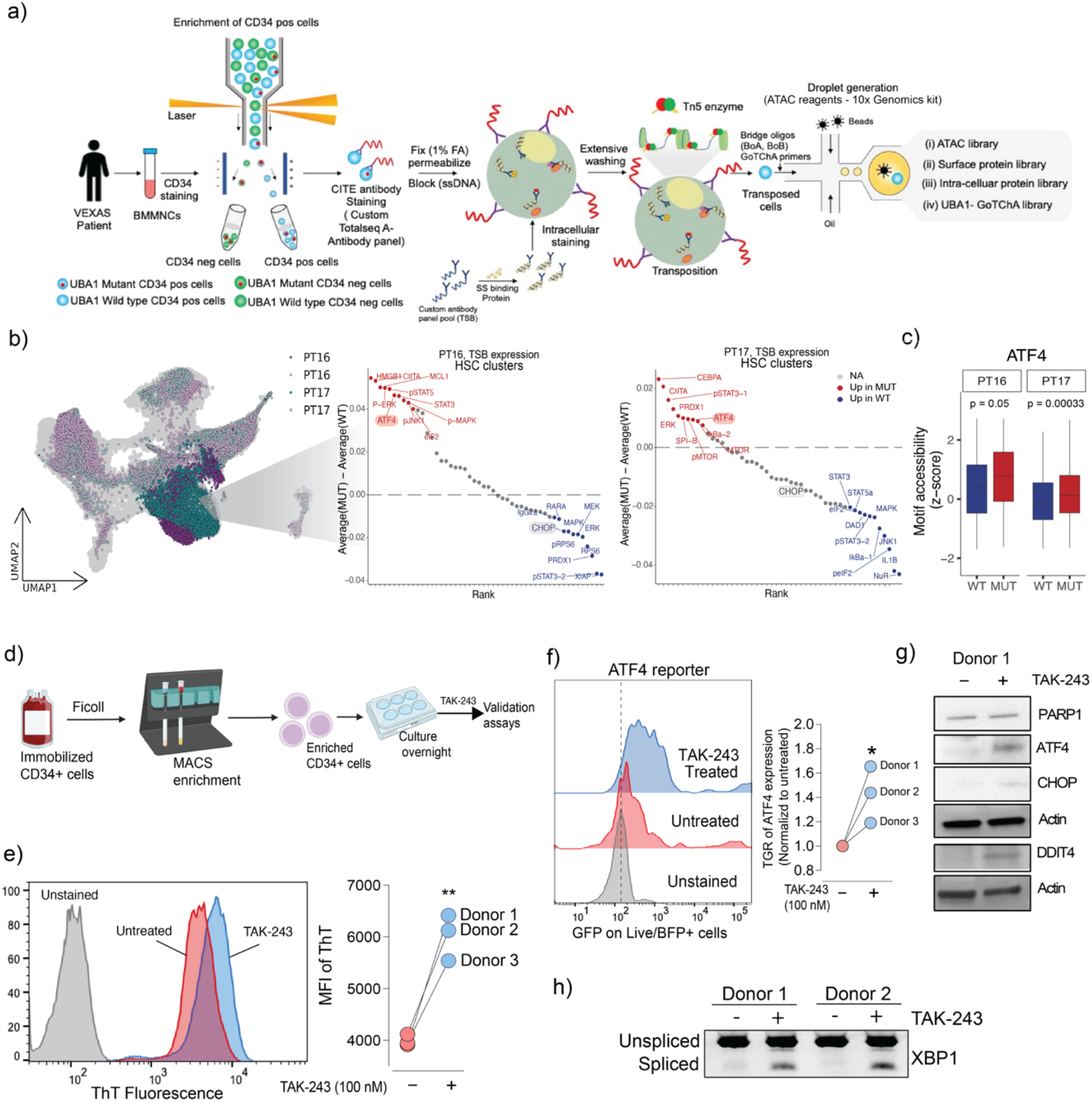
Intracellular protein landscape of CD34+ *UBA1*-mutant cells. **(a)** Schematic for GoT-ChA + Phospho-seq approach for simultaneous profiling of genotype, chromatin accessibility, cell surface proteins and intracellular proteins. **(b)** Left, UMAP for two samples from VEXAS patients constructed using the snATAC-seq data (n = 25,711 cells total). Cells are colored by patient samples. Right, rank of differential intracellular protein expression levels assessed by TSB custom antibody panel (Supplementary Table 13) detection showing those proteins with increased expression levels in mutant (red) and wild-type (blue) HSCs for each patient [PT16 (n = 62 wild-type and n = 146 mutant cells) and PT17 (n = 359 wild-type and n = 532 mutant cells)]. ATF4 and CHOP are highlighted as increased and decreased in both patients, respectively. **(c)** Transcription factor motif accessibility for ATF4 in HSCs from PT16 (n = 62 wild-type and n = 146 mutant cells) and PT17 (n = 359 wild-type and n = 532 mutant cells). Error bars represent the minimum and maximum values as determined by 1.5 × IQR (outliers not shown), boxes represent the interquartile range and lines represent the median. Two-sided Wilcoxon rank sum test. **(d)** Schematic for ex vivo culture system for CD34+ expansion and treatment with UBA1 inhibitor TAK-243 for functional validation assays. **(e)** Thioflavin T (ThT) fluorescence assay to measure proteomic aggregation in CD34+ cells treated with UBA1 inhibitor TAK-243 and untreated cells. Left, FACS profile showing right shift reflecting increased ThT fluorescence in UBA1-inhibited cells. Right, quantification mean fluorescence intensity (MFI) of ThT in treated and untreated cells for 3 donors. ** indicates significance (*P* = 0.001), two-sided Student t-test. (**f**) ATF4 reporter activity in CD34+ cells treated with TAK-243 (100 nM) compared to untreated cells (representative histogram showing ATF4 GFP expression gated on live, TagBFP+ cells). The transgene ratio (TGR) was calculated (MFI of GFP/MFI of BFP) and normalized to untreated cells. (MFI, mean fluorescence intensity, n = 3 donors). * Indicates significance, (*P* = 0.03), two-sided Student t-test. **(g)** Western blot analysis to assess protein levels of PARP1 (control marker to show that UBA1 inhibition ex vivo does not significantly impact cell survival), ATF4, CHOP and DDIT4 in TAK-243-treated and untreated CD34+ cells from Donor 1. Results are representative of n ≥ 3 technical replicates. **(h)** Detection of spliced versus unspliced *XBP1* mRNA in UBA1-inhibited CD43+ cells from two donors.

### PERK-ATF4 signaling is activated in *UBA1*-inhibited CD34+ cells

To functionally validate these findings, we expanded ex vivo mobilized HSCs from discarded allogeneic transplantation grafts obtained from healthy donors, with further *ex vivo* induction of myeloid differentiation (Methods; **Fig. 5d**). We then inhibited UBA1 ex vivo by applying TAK-243, a small molecule inhibitor of UBA1^15^, to our cultured normal CD34+ cells to model the VEXAS phenotype (**Supp. Fig. 13**). We first confirmed that TAK-243 effectively targeted the ubiquitin pathway through analyzing global reduction of ubiquitinated protein (**Supp. Fig. 13a**) which resulted in accumulation of protein aggregation measured via Thioflavin T (ThT) fluorescence as a marker of protein aggregation indicative of protein degradation failure, observing increased ThT signal in treated versus untreated cells across three donors (**Fig. 5e**). Next, to confirm the activation of *ATF4* in the ex vivo system, we performed an *ATF4* reporter assay as previously described^40^ (**Supp. Fig. 14a**). The transgene ratio in CD34 cells treated with TAK-243 at the end of 48 hours showed an increased activation of *ATF4* compared to untreated cells **(Fig. 5f**), suggesting an activation of the UPR pathway via PERK. At the transcript level, *ATF4* as well as downstream ATF4 target *DDIT4*^*50*^ were significantly increased upon UBA1 inhibition, confirming our findings from single-cell analysis in VEXAS patients **(Supp. Fig. 14b)**. These findings were further supported through Western blot analysis that showed an increase of ATF4 and DDIT4 in treated cells, indicating an active PERK response (**Fig. 5g**). Importantly, levels of CHOP do not similarly increase upon TAK-243 treatment, supporting the model that PERK signaling through survival-promoting *ATF4*, and not the pro-apoptotic CHOP pathway, is induced by UBA1 inactivation, potentially underlying the ability of *UBA1*-mutant HSCs to survive in VEXAS. Notably, we also observed increased splicing of *XBP1* RNA in these CD34+ cells treated with TAK-243 inhibitor (**Fig. 5h**), further indicating a co-activation of the IRE1 pathway.

Together, these results confirm *PERK* and *IRE1* activation at the intracellular protein/RNA level in UBA1-inhibited HSPCs, and show that our CD34+ expansion system allowing for UBA1 inhibition ex vivo recapitulates VEXAS phenotypes of *UBA1*-mutant cells, providing a platform to test potential agents for targeting UBA1-inhibited HPSCs.

To confirm our findings implicating ATF4 activation in response to proteomic stress, and taking advantage of our ex vivo VEXAS model system, we profiled ATF4 binding directly using CUT&RUN in TAK-243-treated cells. We called binding peaks from bulk sequencing data, and annotated these peaks for overlap with promoter regions. We performed gene set enrichment analysis of the ATF4-bound promoter peaks, identifying enrichment of ER stress response, XBP1 target genes, ATF4 target gene module, Myc activation and MAPK signaling pathways (**Fig. 6a, Supp. Table 16**). Inspection of individual loci showed increased binding of ATF4 in UBA1-inhibited cells at promoter regions of genes involved in myeloid differentiation (e.g., *SPI1*) and inflammation (e.g., *TNFRSF14*; **Fig. 6b**). These data demonstrate that increased ATF4 levels in UBA1-inhibited cells also reflect increased DNA binding of ATF4 at functionally important genes that likely contribute to the VEXAS phenotype of proteomic stress response, increased myeloid bias and inflammation.

**Figure 6:**
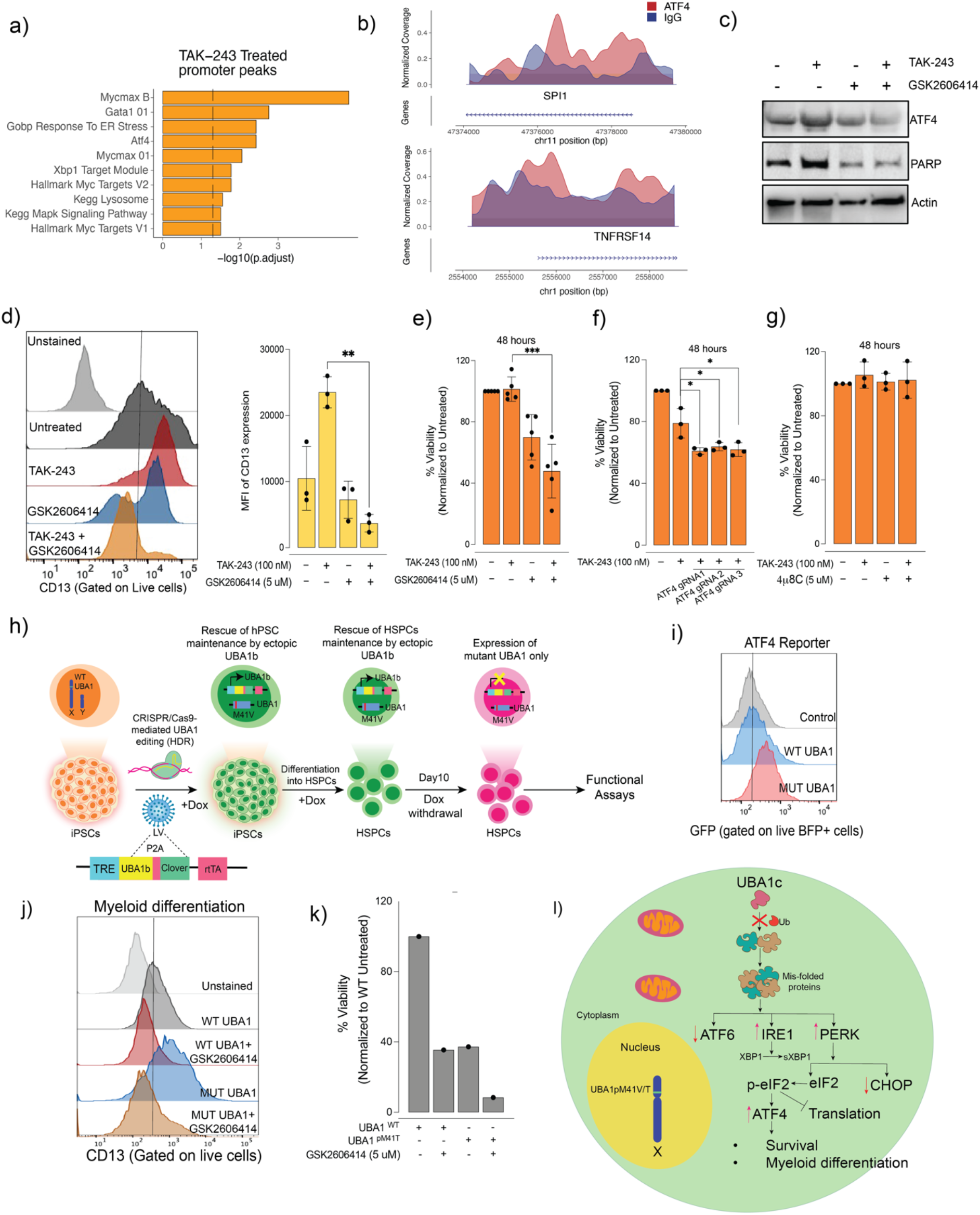
PERK inhibition reduces myeloid bias and survival in VEXAS model HSCs. **(a)** Geneset enrichment analysis of ATF4-bound promoter peaks obtained from CUT&RUN experiments in UBA1-inhibited (TAK-243-treated) CD34+ cells (500,000 live cells). **(b)** Example ATF4 CUT&RUN binding at promoter regions of genes involved in myeloid differentiation (e.g., *SPI1*) and inflammation (e.g., *TNFRSF14*) in UBA1-inhibited (red) cells versus IgG-treated (blue) cells. **(c)** Western blot showing increased expression of ATF4 in CD34+ cells upon UBA1 inhibition (TAK-243 - 100nM) and decreased expression when combined with PERK inhibitor (GSK2606414 - 5uM), correlated with PARP1 expression. Results are representative of n ≥ 3 biological replicates. **(d)** Myeloid bias rescue with co-inhibition of PERK (GSK2606414) and UBA1 (TAK-243). Left, FACS profile of unstained, untreated, UBA1-inhibited, PERK-inhibited or co-inhibited CD34+ cells cultured in myeloid differentiation media (n = 3 donors) ex vivo. Right, quantification of DAPI-/ CD13+ expression in untreated, UBA1-inhibited, PERK-inhibited or co-inhibited CD34+ cells from 3 donors. Error bars represent standard error. ** indicates significance (*P* = 0.001), two-sided Student T test. **(e)** Quantification of cell viability in untreated, UBA1-inhibited (TAK-243 -100 nM), PERK-inhibited (GSK2606414 - 5uM) or co-inhibited (TAK-243 + GSK2606414) CD34+ cells from 5 independent experiments at the end of 48 hours. Viability was assessed by AnnexinV and PI staining. *** indicates significance (P < 0.001); error bars represent standard error. Two-sided Student t-test. **(f)** Quantification of cell viability in Scramble, UBA1-inhibited (TAK-243), *ATF4* knockout CD34+ cells from 3 donors at the end of 48 hours. Viability was assessed by AnnexinV and PI staining. * indicates significance (*P* = 0.03); error bars represent standard error. Two-sided Student t-test. **(g)** Quantification of cell viability in untreated, UBA1-inhibited (TAK-243 - 100nM), IRE1-inhibited (4u8C - 5uM) or co-inhibited (TAK-243 + 4u8C) CD34+ cells from 3 donors at the end of 48 hours. Viability was assessed by AnnexinV and PI staining. *** indicates significance (*P* < 0.001); error bars represent standard error. Two-sided Student t-test. **(h)** Modelling of *UBA1*^*M41V*^ mutation in CD34+ cells derived from iPSCs. **(i)** ATF4 reporter activity in *UBA1*^*M41V*^ CD34+ derived from iPSCs compared to control HSCs cells (histogram showing ATF4 GFP expression gated on live, TagBFP+ cells). **(j)** Myeloid bias rescue with PERK inhibitor (GSK2606414) on *UBA1*^*M41V*^ CD34+ derived from iPSCs compared to control HSCs cells. FACS histogram showing unstained, control HSCs and mutant HSCs along with PERK-inhibition in myeloid differentiation media tested ex vivo. CD13 was used as a marker of myeloid differentiation. **(k)** Quantification of cell viability in control and mutated CD34+ cells treated with PERK inhibitor (GSK2606414 – 5uM) at the end of 48 hours. Viability was assessed by AnnexinV and DAPI staining. **(l)** Model of HSCs in VEXAS. UBA1p.M41V/T somatic variant leads to impaired ubiquitination, driving proteomic stress induced by misfolded proteins. This triggers the unfolded protein response (UPR), with activation of the IRE1-and PERK-mediated pathways, but not the CHOP pathway that shunts cells towards apoptosis. The IRE1 pathway is upregulated, reflected by increased splicing of *XBP1*, but as inhibition of IRE1 in UBA1-inhibited cells does not decrease survival, the IRE1 pathway is not essential for mitigating proteomic stress in VEXAS cells. In contrast, PERK activation leading to increased expression of ATF4 in the context of upregulated inflammation pathways leads to increased HSC survival, with myeloid bias. Inhibition of PERK in UBA1-inhibited HSCs reverses the myeloid differentiation bias and decreases cells survival, suggesting that PERK-mediated UPR sufficiently manages proteomic stress in the context of impaired ubiquitination to rescue cells from apoptosis, leading to increased cell survival.

### PERK inhibition is a novel potential strategy for VEXAS treatment

Given these results nominating *ATF4* induction as a potential survival mechanism in *UBA1*-mutant/inhibited cells, we next tested the effect of PERK pathway inhibition in our ex vivo VEXAS model using the PERK inhibitor GSK2606414^53^. Treating normal CD34+ cells with UBA1 inhibitor led to increased ATF4 levels compared to untreated cells; however, co-inhibition of UBA1 together with PERK inhibition decreased ATF4 levels with concomitant decrease in viability, evidenced through reduced PARP1 expression (**Fig. 6c**), indicating that PERK inhibition effectively downregulates ATF4 signaling and affects survival.

To test the effect of PERK inhibition on myeloid differentiation in our VEXAS model, we assessed myeloid bias following ex vivo differentiation in the presence or absence of UBA1 and PERK inhibitors, measuring expression of myeloid marker CD13^54^ via FACS (Methods). While cells treated with UBA1 inhibitor showed the expected shift reflecting bias towards myeloid differentiation, this pattern was reversed with the addition of PERK inhibitor, which restored unbiased differentiation in cells treated with both inhibitors (**Fig. 6d**).

Previous studies^55^ showed that PERK induces apoptosis as part of the UPR when activated via CHOP, but in our VEXAS model, PERK is protecting the cells. We speculated that the activation of pro-survival versus pro-apoptotic pathways might be driven by a low level of misfolded protein accumulation in HSCs, as high levels of proteotoxic stress that induce a strong UPR would favor apoptosis^40^. To test this hypothesis, we treated our UBA1-inhibited CD34+ cells with tunicamycin, a strong inducer of UPR, to increase the level of proteotoxic stress in these cells. We observed decreased survival in the cells treated with both the UBA1 inhibitor and UPR inducer (**Supp. Fig. 15a**), suggesting that the UBA1 inhibitor induces lower levels of misfolded protein stress, activating the PERK pathway to predominantly potentiate an anti-apoptosis program. Notably, adding PERK inhibitor to tunicamycin rescued the cells from tunicamycin-induced apoptosis (**Supp. Fig. 15b**), which also has been reported previously^40^, further supporting the pro-apoptotic function of PERK signaling in the context of higher levels of proteotoxic stress. Thus, inhibiting PERK by GSK2606414 or AMG PERK44 in the presence of TAK-243 or PYR-41 (UBA1 inhibition) induces higher levels of cell killing (**Fig. 6e, Supp. Fig. 15c**,**d**), suggesting that PERK is induced as a rescue mechanism in the context of more moderate proteotoxic stress.

To further validate that PERK is leading to increased survival in UBA1-inhibited cells specifically through ATF4 activation, we knocked out *ATF4* in CD34+ cells. Following treatment with UBA1 inhibitor, we observed that in the absence of *ATF4* (**Supp. Fig. 15e**), UBA1 inhibition significantly affected the viability at the end of 48 hours (**Fig. 6f**). Finally, we assessed cell viability of UBA1-inhibited cells treated with either PERK inhibitor or an IRE1 inhibitor (4μ8c^56^) to assess the contribution of these different UPR pathways to cell survival in the context of reduced ubiquitination. Notably, we observed that dual inhibition of UBA1 together with PERK reduced cell survival. In contrast, co-inhibition of UBA1 and IRE1 did not affect the viability of these HSPCs (**Fig. 6g**), suggesting that IRE1 activation is not required for maintaining VEXAS HSC survival.

As UBA1 inhibition may not fully recapitulate the mutant phenotype, we further generated isogenic induced pluripotent stem cells (iPSCs) with the *UBA1*^*M41V*^ mutation through CRISPR/Cas9-mediated gene editing (**Fig. 6h** and **Supp. Fig.16a**,**b**). Despite efficient gene editing, iPSCs with the *UBA1*^*M41V*^ mutation were rapidly lost upon passaging and could not be expanded. This negative selection specifically of M41-mutant clones over clones with different editing outcomes strongly suggested that the *UBA1*^*M41V*^ mutation was incompatible with iPSC maintenance. To overcome this limitation, we combined gene editing with complementation of UBA1b expression (isoform sufficient for ubiquitination) through a doxycycline (dox)-inducible lentiviral vector (**Fig. 6h** and **Supp. Fig. 16c-e**). This allowed for selection of several single iPSC clones with the *UBA1*^*M41V*^ mutation to be expanded and maintained in the presence of Dox. Upon hematopoietic differentiation into HSPCs and Dox withdrawal, HSPCs that lacked expression of the UBA1b isoform instead expressed the UBA1c isoform, as seen in mutant cells from VEXAS patients (**Supp. Fig. 16f,g**). Using this model, we observed that *UBA1*^*M41V*^-mutant iPSC-derived HSPCs showed increased expression of ATF4 (**Fig. 6i**) and myeloid bias (**Fig. 6j**), compared to isogenic wild-type HSPCs (parental line), and PERK inhibition reversed these expression patterns. In addition, *UBA1*^*M41V*-^mutant cells were more sensitive to PERK inhibitors than the isogenic wild-type cells (**Fig. 6k**). Collectively, our VEXAS ex vivo model mechanistically validates that PERK contributes to the survival of *UBA1*-mutated HSCs via ATF4 activation (**Fig. 6l**).

## Discussion

VEXAS syndrome presents a unique opportunity to study how somatic mutation perturbs cellular pathways and hematopoietic differentiation to lead to inflammatory disease. Here, using genotype-aware single-cell multi-omics applied directly to patient samples, we characterized the consequences of *UBA1* mutation in disease-initiating HSCs, identifying pro-survival mechanisms that have the potential to be exploited for therapeutic targeting. As current VEXAS treatment options are focused on curtailing inflammation, our findings present new opportunities for effective intervention strategies designed to eliminate disease-initiating HSCs for clonal eradication.

Genotype-aware single-cell multi-omics has emerged as a powerful tool for dissecting the cellular consequences of somatic mutations in human blood disorders, where single-cell genotype-to-phenotype mapping elucidated mechanisms underlying clonal expansions driven by DNA methylase^24^ mutations in CHIP or splicing factor mutations in MDS and CHIP^20^. Importantly, this approach removes confounding from inter-patient variation and identifies the cell-intrinsic effects of somatic mutations across different cell types. Harnessing this framework, we applied a suite of multi-omics tools to comprehensively characterize mutant versus wild-type cells from VEXAS patients across molecular layers, including transcriptome, chromatin, cell-surface and intracellular proteins.

Through this analysis, we assessed *UBA1* mutation enrichment across cell types, finding the previously reported myeloid enrichment along with the unexpected observation of mutation enrichment in NK cells, despite overall depletion of mutants in lymphoid populations. Two potential explanations for the unexpected divergence between NK cells and other lymphoid cells in mutant cell enrichment include a possible shared ontogeny between myeloid and NK cells^57^, or shared selective pressures encountered by mature monocytes and NK cells that favor mutated cells. Indeed, the presence of myeloid-biased mutations (*TET2*) in NK cells was previously reported in the context of MDS^58^ and CHIP^59^. Notably, mutant NK cells showed increased expression of inflammatory and cytotoxic pathways, suggesting enhanced activity. We therefore speculate that the distinct phenotypes observed in *UBA1*-mutant NK cells may also contribute to the manifestation of inflammatory disease, which in previous analysis of VEXAS patients was primarily ascribed to mutated monocytes. How these mutant NK cells impact VEXAS pathology or can be exploited as a therapeutic target are exciting avenues for future exploration.

A major advantage of the single-cell genotype-phenotype mapping is the ability to identify pathways that were differentially activated specifically in *UBA1*-mutant HSCs. The identification of mutation-specific vulnerabilities has the potential to lead to the development of targeting strategies that specifically eliminate *UBA1*-mutant progenitor cells while sparing wild-type HSCs. While recent work used scRNA-seq to capture genotype information from 10x sequencing data from VEXAS HSPCs^17^, the total cells analyzed and the genotyping efficiency were low, particularly for HSCs (199 HSCs genotyped). In comparison, our GoT approach with targeted amplification enables high-confidence genotyping calls, allowing for enhanced analysis of the HSC population that is under-represented in CD34+ cells (3,211 HSCs genotyped). We also observed that HSPCs had a higher genotyping efficiency compared to total BMMNCs. This increased genotyping efficiency, combined with the profiling of additional molecular modalities, uncovered the cell-intrinsic signatures of *UBA1* mutation in VEXAS HSCs, including increased inflammation, myeloid bias and activation of the *ATF4* arm of the PERK UPR pathway. Notably, we observed that mutant cells had increased *ATF4* levels at both the transcriptomic and intracellular protein levels, as well as increased accessibility at the promoters of ATF4-target genes. We validated these results ex vivo using pharmacological inhibition of UBA1 in CD34+ cells, providing functional evidence that *ATF4* is activated when UBA1 is disrupted, further suggesting that the PERK pathway is an important component of the UPR in VEXAS HSCs.

As inhibitor-mediated UBA1 inhibition of CD34+ cells may not fully recapitulate the mutant phenotype, we generated isogenic iPSCs with the *UBA1*^*M41V*^ mutation through CRISPR/Cas9-mediated gene editing. Notably, like many cellular models, these iPSCs were negatively impacted by *UBA1*^*M41V*^ mutation, which motivated us to create an alternative strategy through introduction of wild-type *UBA1* under doxycycline in these edited iPSCs. Through this system we were able to maintain the *UBA1*^*M41V*^ mutation and differentiate cells into HSPCs. Upon the removal of doxycycline, wild-type *UBA1* was inhibited, allowing for the expression of only the *UBA1*^*M41V*^ mutation in these HSPCs. We took advantage of this model to validate our findings, showing that the PERK branch of UPR is activated and influences myeloid differentiation. We also showed that these mutant cells are more sensitive to PERK inhibition than the wild-type cells, validating our proposed mechanism for enhanced survival of mutated cells through proteotoxic stress.

The UPR cascade is complex and versatile, adapting to cellular needs depending on the level and type of proteomic stress and overall state of the cell. HSCs are sensitive to protein stress^40^, with UPR activation during stress conditions playing an important role in maintaining the integrity of the HSC pool^60^. While PERK has been reported to induce apoptosis when activated via CHOP, in our study we consistently observed no induction of this pro-apoptotic CHOP pathway across modalities in *UBA1*-mutant HSCs. Our ex vivo experiments show that with high proteotoxic stress as with tunicamycin, PERK is pro-apoptotic and its inhibition is protective. In contrast, in modest proteotoxic stress, such as with UBA1 inhibition or mutation, PERK is pro-survival and its inhibition leads to reduced HSC viability. Altogether, our data suggest that PERK activation underlies the enhanced survival of *UBA1*-mutant HSCs through mounting a survival-promoting response to proteotoxic stress that avoids apoptosis. Excitingly, this nominates a strategy for targeted elimination of mutant HSCs in VEXAS, as PERK inhibitors have been characterized in preclinical models^61^ and are currently being tested in clinical trials for other indications (NCT04834778).

Eliminating myeloid-biased HSCs has been shown to recover lympho-biased and balanced HSCs in mice, reducing immuno-aging phenotypes^62^. Previous studies on CHIP have demonstrated myeloid bias of HSCs influenced by *TET2* mutation^63^, but the molecular mechanisms that govern such bias are not clear. In this study, we show that inhibiting PERK along with UBA1 resulted in significant reduction of myeloid output of HSCs. PERK has already been shown to play a major role in myeloid commitment of HSCs^64^. Moreover, we demonstrate that PERK downstream transcription factor, ATF4, also known to contribute to myeloid differentiation^65^, is upregulated in VEXAS HSCs and binds at myeloid genes. Thus, through this study, we have established a strong link between PERK activation and myeloid bias in *UBA1*-mutated HSCs. Whether aged HSCs with CHIP mutations also have default activation of the PERK-ATF4 pathway is a future question of interest, potentially opening up new ways to target CHIP clones to eliminate myeloid-biased HSCs.

Collectively, single-cell genotype to phenotype mapping of VEXAS syndrome revealed cell-intrinsic differences in somatically mutated cells that contribute to HSC survival. Thus, our analyses demonstrate the power of single-cell multi-omics profiling of somatic mosaicism in primary patient samples for discovery of disease mechanisms as well as novel candidates for precision targeting of mutant cells.

## Acknowledgments

We thank the members of the Landau laboratory and for their advice and discussions. Sa.G. is supported by the American Society for Hematology Fellow-to-Faculty Scholar Award. E.O.E. is supported by a Medical Scientist Training Program grant from the National Institute of General Medical Sciences of the National Institutes of Health under award number: T32GM007739 to the Weill Cornell/Rockefeller/Sloan Kettering Tri-Institutional MD-PhD Program. F.I. is supported by the American Society of Hematology Fellow-to-Faculty Scholar Award number 204377-01. J.D.B. is a postdoctoral fellow of the Jane Coffin Childs Memorial Fund for Medical Research. D.B.B. is supported by Gabrielle’s Angel Foundation, Department of Defense (BMFRP-IDA HT9425-23-1-0507), Edward P. Evans Foundation and the AAMDSIF. D.A.L. is supported by the Burroughs Wellcome Fund Career Award for Medical Scientists, Valle Scholar Award, Leukemia Lymphoma Scholar Award and the Mark Foundation Emerging Leader Award. This work was also supported by the Tri-Institutional Stem Cell Initiative, the National Cancer Institute (R33 CA267219), the National Heart Lung and Blood Institute (R01HL157387-01A1), the National Human Genome Research Institute Center of Excellence in Genomic Science (RM1HG011014) and the National Institutes of Health Common Fund Somatic Mosaicism Across Human Tissues (UG3NS132139-01). This work was made possible by the MacMillan Family Foundation and the MacMillan Center for the Study of the Non-Coding Cancer Genome at the New York Genome Center.

## Author Contributions

SaG, PS, OK and DAL conceived the project. SaG, DBB and DAL devised the research strategy. SaG, RMM, JS, EOE, TB, KB, ACI, NC, CC, FI, ZW, ShG, NSY, JDB, RS, BT, MH, IR, EPP, PS, OK, DBB and DAL analyzed the data. SaG, JS, KB, ACI performed experiments. SaG, RMM, JS, EOE, KT, FI, ZW, ShG, NSY, JDB, RS, BT, MH, IR, EPP, PS, OK, DBB and DAL helped interpret results. SaG, RMM, CP, EPP and DAL wrote the manuscript. RMM, EOE, TB, NC and CC performed computational analysis. KT generated the VEXAS iPSC lines. FI advised on GoT-ChA analysis. SM assisted in generating the iPSC *UBA1*-mutant cells. ZW, ShG, NSY, BT, MH, PS, OK and DBB provided clinical samples. JDB and RS advised on Phospho-seq analysis. EPP devised the research strategy to generate the iPSC lines. All authors reviewed and approved the final manuscript.

## Competing Interests

DBB has served as a consultant for Sobi, Novartis and Alexion Pharmaceuticals. KT is an employee of Daiichi Sankyo Co., Ltd. EPP has received consulting fees or honoraria from Janssen and Cellarity, unrelated to the current study. DAL has served as a consultant for Abbvie, AstraZeneca and Illumina and is on the Scientific Advisory Board of Mission Bio, Pangea, Alethiomics, and C2i Genomics; and has also received prior research funding from BMS, 10x Genomics, Ultima Genomics, and Illumina unrelated to the current manuscript.

## Data and Code Availability

The single-cell multimodal sequencing data reported in this paper will be deposited in GEO and EGA. Raw data and processed data files from the samples reported in Wu et al. are available at GEO as part of superseries GSE196052. The GRCh38 reference genome was used for alignment of single cell RNA-seq data (refdata-gex-GRCh38-2020-A) and for single cell ATAC-seq data (refdata-cellranger-atac-GRCh38-1.2.0), freely available from the 10x Genomics website (https://support.10xgenomics.com). All analysis scripts can be found https://github.com/RebeccaMurray/VEXAS-scRNA-seq https://github.com/RebeccaMurray/VEXAS-scATAC-seq. at and

## Methods

### Human VEXAS samples

The study was approved by the local ethics committee and by the Institutional Review Board of New York University, Weill Cornell Medicine, Cochin Institute (Paris, France), and Hospices Civils de Lyon (France) conducted in accordance with the Declaration of Helsinki protocol. All patients provided informed consent. Cryo-preserved bone marrow mononuclear cells or peripheral blood mononuclear cells from patients with documented *UBA1* mutations were retrieved. Cryopreserved bone marrow mononuclear cells were thawed and stained using standard procedures (10 min, 4°C) with the surface antibody CD34-PE-Vio770 (clone AC136, lot no. 5180718070, dilution 1:50, Miltenyi Biotec) and DAPI (Sigma-Aldrich). Cells were then sorted for DAPI−, CD34+ and DAPI−, CD34− cells using BD Influx at the Weill Cornell Medicine flow cytometry core. No statistical methods were used to predetermine sample size. The experiments were not randomized and investigators were not blinded to allocation during experiments and outcome assessment.

#### Genotyping of Transcriptome with CITE-seq

GoT-CITE-seq was performed according to our previous publications (GoT^16^, GoT-Splice^18^). Briefly, live cells after sorting were stained for surface antibodies tagged with barcoded oligos (TotalSeq™-C Human Universal Cocktail, V1.0, 399905, 1 vial per 0.5 million cells), washed and subjected to scRNA-seq using standard 10x Genomics Chromium 5′ library kit according to the manufacturer’s recommendations with following modifications. a) The surface receptor protein library was generated from the supernatant saved during the full-length cDNA clean-up step as previously described^25^. b) For the UBA1 GoT library, 10% of the full-length cDNA was used to generate the UBA1 GoT library and XBP1 GoT library using *UBA1* and *XBP1*-specific primers (**Supp. Table 17** and **Supp. Fig. 1**).

### ATAC with Select Antigen Profiling by sequencing (ASAP-seq) with GoT-ChA

ASAP-seq with GoT-ChA was performed as described previously^19^. Briefly, sorted cells are stained for surface receptor antibodies tagged with barcoded oligos (TotalSeq™-A Human Universal Cocktail, V1.0), washed, fixed (1% formaldehyde), permeabilized and subjected to transposition of genomic DNA as per 10x Genomics Chromium ATAC-seq kit instruction. During the cell barcoding amplification reaction, primers to capture *UBA1* mutation (GoT-ChA primers - 22 uM) and bridging oligos to capture the antibody tag are added. The sample is then split, with 10% used for genotyping and the remaining 90% used for assay for transposase-accessible chromatin (ATAC) library construction. The supernatant from the first purification is used to generate the surface protein libraries. The genotyping, single-cell ATAC and single-cell protein libraries are then intersected via shared barcodes. *UBA1*-specific primers for GoT-ChA are provided in **Supp Table 17**.

### Phospho-seq with GoT-ChA

Phospho-seq with GoT-ChA was done according to the previous publication^32^ along with the following modifications. Briefly, sorted cells are fixed, permeabilized, blocked and stained with intracellular antibodies tagged with barcoded oligos (**Supp. Fig. 11d;** custom made - antibody list in **Supp. Table 14**,**18**) subjected to transposition of genomic DNA. During the cell barcoding amplification reaction, GoT-ChA primers to capture *UBA1* mutation and bridging oligos to capture the antibody tag are added. The sample is then split, with 10% used for genotyping and the remaining 90% used for assay for transposase-accessible chromatin (ATAC) library construction. The supernatant from the first purification is used to generate the intra-cellular protein libraries. The genotyping, single-cell ATAC and single-cell protein libraries are then intersected via shared barcodes.

### Mobilized CD34+ HSPCs cultures

Human CD34+ HSPCs from mobilized peripheral blood of healthy adults (discarded allogeneic transplants) were obtained from Weill Cornell Medicine, Blood bank, NYC, USA. The cells were thawed and cultured in StemSpan SFEM II culture medium, supplemented with 1X CC100 (containing the cytokines FLT3L, SCF, IL3, and IL6), recombinant thrombopoietin (TPO) (100 ng/ml), and UM171 (35 nM), as described previously^66^. The cells were treated with drugs [UBA1 inhibitor TAK-243 (100 nM) and PYR-41 (5 uM) PERK inhibitors GSK2606414 (5 uM) and AMG PERK-44 (1 uM); and IRE1 inhibitor 4μ8C (5 uM)] for the time points indicated for each experiment.

### Myeloid differentiation

The CD34 cells were cultured in StemSpan SFEM II culture medium, supplemented with 1X StemSpan™ Myeloid Expansion Supplement (Stemcell Technologies, Catalog # 02693) containing SCF, TPO, G-CSF and GM-CSF. The cells were cultured for 7 days (with and without drugs) in this media and at the end of 7 days, the myeloid differentiation potential of CD34 was assessed by measuring the expression of CD13 on these cells by flow cytometry (FACS Canto, BD).

### XBP1 splicing

CD34+ cells after TAK-243 drug treatment were collected, RNA was isolated and cDNA was produced using iScript cDNA Synthesis Kit (Bio-Rad). Total *XBP1* was amplified via PCR using *XBP1*-specific primers (**Supp. Table 17**). The PCR products (unspliced 164 bp and spliced 138 bp) were visualized on a 4% agarose gel containing SYBRsafe.

### Measurement of intracellular mis-folded protein aggregates

The measurement of intracellular accumulation of mis-folded protein aggregates was measured using ThioflavinT assay as described^67^. Briefly, the CD34+ cells post treatment with UBA1 inhibitor were spun down, washed with PBS and stained for thioflavin-T (ThT) for 15 min in PBS at room temperature in the dark, followed by washing with PBS and staining with live dead stain (eBioscience™ Fixable Viability Dye eFluor™ 780) for 5 min. The cells were washed and acquired in flow cytometry (FACS Canto, BD) and the data were analyzed using FlowJo software.

### Apoptosis Assay

The CD34+ cells were isolated, cultured as described above and treated with the inhibitors for 48 hours. After 48 h of incubation at 37 °C in a CO_2_ incubator, the CD34+ cells were carefully pipetted out and their viability was measured using Annexin V apoptosis assay kit (BD Pharmingen, San Diego, CA, USA) as per the manufacturer’s protocol and counter stained with DAPI.

### Lentiviral transduction and ATF4 reporter assay

ATF4 reporter Viral particles were made from pSMALB-ATF4.5 vector (a gift from John Dick & Peter van Galen; Addgene plasmid # 155032; http://n2t.net/addgene:155032; RRID:Addgene_155032) pseudotyped with the vesicular stomatitis virus G (VSVG) protein using the pMD.G vector, and psPAX2 was used for packaging in 293T cells using FuGENE® HD Transfection Reagent (Promega, cat no. E2311). Lentivirus was concentrated using Lenti-X Concentrator (Takara Bio cat no. 631232), resuspended in StemSpan SFEM II supplemented with 1% BSA and stored at –80 °C until use. Cord blood cells were thawed, plated and infected with lentiviral suspension in a total volume of 100 μl (96 well plate). After 16 h, fresh media was added to expand the cells. After 24 h the cells were treated and the readings were taken at the end of 48 h post treatment. The transgene ratio between GFP and TagBFP was calculated as previously reported^40,60^.

### Cas9 protein electroporation mediated knockout

The knockout of *ATF4* was achieved using the electroporation method; the gRNAs targeting the *ATF4* gene were ordered from IDT. Briefly, 1.2 ul of 100 mM each gRNA was incubated with 2.1 ul of 62 mM Alt-R™ S.p. Cas9-GFP V3 ribonuclear protein (IDT, 10008161) for 20 min at room temperature. Following the CD34+ cell wash with DPBS, cells were resuspended in 20 uL of P3 Lonza buffer with supplement (Lonza, V4XP-3032). 5 uL of the Cas9 ribonucleoprotein master mix and 1 uL of 100 mM Electroporation enhancer (IDT) were added to the cells, gently mixed three times, and transferred to an electroporation cuvette. Cells were electroporated using the DZ-100 program in a 4D-Nucleofector X Unit (20 uL cuvettes). Immediately after electroporation, 80 uL of prewarmed media were added to the electroporation cuvette, which was placed in an incubator at 37°C for 5 min. Cells were then plated at a density of 500,000 cells/mL in the complete media. On the next day, the media was changed and cells were treated with drugs for 48 h. At the end of incubation, a small aliquot of cells was taken for RNA extraction and Q-PCR was carried out to calculate the efficiency of knockout.

### Generation of a human iPSC model of VEXAS syndrome

We used the previously described normal iPSC line N-2.12-D-1-1 as the parental line^68^. We used CRISPR/Cas9-mediated homology-directed repair (HDR) to introduce the *UBA1*^*M41V*^ mutation using nucleofection of RNP with gRNA and M41V mutant donor DNA (see **Supp. Fig. 16**). A lentiviral vector expressing the UBA1b cDNA linked to Clover fluorescent protein through a P2A peptide under control of the tetracycline responsive element (TRE) and rtTA was constructed and packaged as described^68^. Culture of human iPSCs on mitotically inactivated mouse embryonic fibroblasts was performed as previously described^69^. All edited *UBA1*^*M41V*^ iPSC lines were maintained in the presence of 1ug/ml of doxycycline (Dox). Hematopoietic differentiation was performed using a spin-EB protocol previously described^69^. Doxycycline was removed from the media on day 10 of differentiation and continued with downstream assays.

### Immunoblot

The CD34 cells after treatment were harvested and the pellets were lysed in RIPA buffer (Sigma, St.Louis, MO, USA) for 30 min on ice, with complete protease inhibitors (Roche, Basel, Switzerland). The lysates were collected by centrifugation at 13,000 rpm for 10 min. The lysates were analyzed in SDS–PAGE. After protein transfer to a nitrocellulose membrane (BioRad, CA, USA), membranes were blocked with non-fat dry milk (5%, 2 h at room temperature) followed by incubation with primary antibodies (**Supp. Table 18**) overnight at 4°C. The blots were washed and incubated with secondary antibodies (Thermo Scientific, IL, USA). The protein bands were detected by the standard chemiluminescence method (Thermo Pierce Femto, Rockford, IL, USA).

### Q-PCR

Total RNA was extracted using Trizol reagent (Invitrogen Carlsbad, CA, USA). 500 ng of the extracted RNA was converted into cDNA using a superscript III cDNA kit (Invitrogen Carlsbad, CA, USA). The expression of genes was studied using the Taqman universal mastermix (Applied Biosystems, Thermo Fischer Scientific, USA) (primer/probe used are provided in **Supp. Table 17**). The Ct values were normalized with *GAPDH* and the fold differences were calculated using the 2^-ΔΔCt^ method.

### CUT&RUN experiments

The CD34+ cells were treated with TAK-243 for 48 h and the cells were collected and immobilized on magnetic beads conjugated with a specific antibody against ATF4 (Cell signaling; (D4B8) Rabbit mAb #11815) as per the instructions from the CUTANA™ ChIC/CUT&RUN Kit (EpiCypher 14-1048). Following overnight incubation with the ATF4/IgG control antibody, the cell-bead mixture was incubated with the pA-MNase fusion protein. The activation of MNase cleavage was triggered by the addition of calcium ions and incubated at 4°C for 2 h, facilitating the selective digestion of DNA associated with ATF4. Released DNA fragments were then collected from the supernatant through a centrifugation step and purified using the column extraction method. For sequencing library preparation, the purified DNA was processed using the CUTANA™ CUT&RUN Library Prep Kit with Primer Set 2 (EpiCypher 14-1002). The workflow included end repair, A-tailing, adapter ligation, and amplification steps using the supplied Primer Set 2. The resulting library was then sequenced on NovaSeq 6000 Illumina machine using the following parameters defined by the kit (paired end - 2 × 50 cycles).

### CUT&RUN analysis

Fastqs were processed by using bowtie2 (v2.5.2) and samtools (v1.9) to align reads to hg38 and then sort reads by position. Reads were then filtered for a minimum mapping quality of 30 and the picard tool MarkDuplicates (v2.27.4) was used to filter out duplicate reads. Peak calling was then performed using macs2 callpeak (v2.2.7.1) with the parameters --nomodel --bw 300 -m 10 30 -- nolambda --extsize 200 -f BEDPE -g hs. A control sample targeting IgG was processed using the same steps and provided as the control to macs2 callpeak using the -c parameter. Peaks were then imported into R and annotated using the annotatr package^70^ (v1.24.0) with hg38 gene promoter regions. The enrichr tool from the clusterProfiler package (v4.6.0) was then used to perform hypergeometric significance testing followed by Benjamini-Hochberg correction using the same set of gene modules used in the RNA and ATAC data GSEA. Coverage plots were generated using the trackplot package^71^.

### scRNA-seq analysis

10x Illumina data were processed using Cell Ranger (v4.0.0) with default parameters and reads were aligned to the human reference sequence GRCh38. All samples were loaded via the Seurat package (v.4.9.9), then QC filtered to remove cells with > 10% mitochondrial reads, < 250 unique genes or < 1,000 Unique Molecular Identifiers (UMIs), or +/-3 s.d. above the median unique genes or UMIs. Doublets were identified using DoubletFinder (v2.0.3) (run with slightly more permissive QC settings, > 500 UMIs and < 20% mitochondrial reads, and estimating a multiplet rate of 0.8% for every 1,000 cells recovered) and removed. SoupX (v1.6.2, run with R v4.0.5) was run using default parameters on the cellranger outputs of each sample to remove ambient mRNA contamination. Samples originally derived from the same donor were merged and then samples were normalized using SCTransform regressing out percent mitochondrial reads and cell cycle (S.Score and G2M.Score). SCTransform also identified the top 3,000 variable genes found in each dataset that are used for integration. Samples were integrated via the FindIntegrationAnchors and IntegrateData functions with k.anchor = 2, k.filter = 5, and k.score = 30, and k.weight = 100 using the rPCA method with the top 50 principal components for each sample. Next, a principal component analysis (PCA) was performed using the variable genes of the integrated dataset, and based on inspection of the elbow plot, 50 PCs were retained for the UMAP algorithm for cluster visualization. Clustering was performed with the FindNeighbors (also using 50 PCs) and the FindClusters (resolution = 2) functions, which rely on the k-nearest neighbors (KNN) algorithm to identify cell clusters. Unique clusters were manually assigned based on the majority predicted cell type labels following label transfer from the Azimuth bone marrow sc-RNAseq reference. Cluster labels were then manually adjusted by differentially expressed genes identified with the FindAllMarkers function which looked only at genes found in at least 25% of cells in either of the two input comparison groups and only returned results for genes with at least a 0.25 log transformed fold change between groups. More specifically, cluster annotations were made according to the differential expression of canonical lineage marker genes identified in previous single-cell RNA-seq data of normal hematopoietic progenitor cells^37^. Cell types were confirmed via surface protein quantification of ADTs.

### IronThrone GoT for processing targeted amplicon sequences and performing mutation calling

Genotyping of single cells was carried out with the IronThrone (v.2.1) pipeline as previously described^16,18^. In brief, individual amplicon reads were assessed for the appropriate structure (*i*.*e*., presence of the primer sequence and the expected sequence between the primer and given mutation site) and all reads were assessed for a matching cell barcode to the list generated from the 10x paired GEX dataset. A Levenshtein distance of 0.1 was allowed for all sequence matching and collapsing steps, and only UMIs with a minimum of 2 supporting reads were retained for final genotyping. Following UMI collapse, genotype assignment of individual UMIs was conducted as described previously with majority rule of supporting reads for wild type or mutant status (using a 0.5 PCR read ratio, above which must be the majority of PCR reads must be for a UMI to be called definitively). Rare UMIs that did not pass this threshold were removed as ambiguous. Additionally, to remove reads that result from PCR recombination, UMIs in the amplicon library that match UMIs of non-*UBA1* genes in the gene expression library were discarded (as described in the IronThrone GoT pipeline^16,18^). The UMIs that did not match any genes in the 10x library were filtered by setting a read-per-UMI threshold at the local minimum of the read count distribution. Given the hemizygous nature of the *UBA1* mutation in male patients, cell barcodes with a minimum of 1 mutant read were genotyped MUT, while those with a minimum of 1 wild-type read were genotyped WT.

### Differential gene expression testing and gene set enrichment analysis

Differential gene expression analysis was performed between mutated and wild-type cells within each progenitor cell type as well as CD14 monocytes. For each cell type, the corresponding cluster was isolated, and patients with fewer than 5 MUT or fewer than 5 WT cells excluded from analysis (all cell types retained at least 3 patients). Mitochondrial and ribosomal genes were removed from the gene count matrix prior to analysis, and then log normalization was run with default parameters for Seurat’s NormalizeData function (scale.factor = 10000, normalization.method = “LogNormalize”, margin = 1). For the progenitor cell types (HSC, EMP, LMPP, GMP, and MkP), genes that are expressed in fewer than 5% of the genotyped cells for each cell type were excluded from analysis. For the CD14 monocytes, cells with nonzero expression of CD3 or CD19 were excluded, and then genes that are expressed in fewer than 10 genotyped monocyte cells were excluded. Linear mixture models were used as previously described^23^ to determine significantly differentially expressed genes while controlling for patient-specific effects.

Gene set enrichment analysis was performed using a custom list of 95 gene modules (**Supp. Table 3**,**7**). Gene modules were obtained from publicly available resources in msigdb via the msigdbr package (v7.5.1) as well as from previous publications^19,72^. For the Han et al. gene sets, conversion from mouse to human genes was performed using the g:Orth tool (https://biit.cs.ut.ee/gprofiler/orth) as previously reported^40,60^.

The fgsea package (v1.24.0) was then used to perform ranked gene set enrichment analysis. Genes were ranked by effect size (avg_log2FC), and p-values were Benjamini-Hochberg corrected for multiple hypothesis testing.

### Gene module score analysis

HSCs were scored for enrichment of the GOBP ATF6, GOBP IRE1, ATF4-only targets (Han et al.) and CHOP-only targets (Han et al.) gene modules using the AddModuleScore function from Seurat (subset of gene sets listed in **Supp. Table 3, 7**). Briefly, the function infers the gene set activity by calculating the average expression levels of each gene set on a single-cell level, subtracted by the aggregated expression of control feature sets. All analyzed features are binned based on averaged expression, and the control features are randomly selected from each bin. Default parameters were used. Distribution of module scores were compared between MUT and WT HSCs and a two-sided Wilcoxon rank sum test was used to determine significance.

### XBP1 splicing analysis

The XBP1 amplicon libraries were processed using kallisto (v0.48.0) and bustools (v0.41.0) via the kb_python package (0.27.2) to quantify UMI counts of the spliced and unspliced isoforms for each cell barcode. These counts were then merged with the scRNA-seq data via shared cell barcodes. To compare *XBP1* splicing rates between mutant and wild-type HSCs, genotyped cells within the HSC cluster were isolated. Cells were then pseduobulked by genotype and the number of spliced and unspliced UMIs counted for both groups. A contingency table was created with rows representing genotype and columns representing unspliced and spliced UMI counts, and then odds ratio and p-value were determined by using a Fisher’s exact test. To compare *XBP1* splicing in HSPCs more broadly, the same test was repeated but including all genotyped cells in the HSC, EMP, LMPP, and MkP clusters. As an alternative way to compare splicing differences in mutant and wild-type HSCs, a sample-aware permutation test was also performed. First, genotyped HSCs were isolated. For each of 10,000 iterations, genotype labels were randomly scrambled within each donor, and then cells were pseudobulked by genotype, the ratio of XBP1s/XBP1u UMIs was calculated for both genotypes, and the difference of (XBP1s/XBP1u)MUT-(XBP1s/XBP1u)WT recorded. This distribution was then compared with the same test statistic (pseudobulking by genotype and calculating (XBP1s/XBP1u)MUT-(XBP1s/XBP1u)WT) obtained with observed genotyping labels, resulting in *P* = 0.0048.

### NK cell analysis

To confidently annotate NK cells, the clusters annotated as T cells or NK cells during the initial cell type annotations were isolated. NKs were then based on the following criteria for RNA expression: CD3D-, CD3G-, CD3E+or-, NCAM-1+ and/or KLRF+, CD14-as reported before^73^. The resultant cells were used in the downstream NK cell analysis (**Fig. 2j,k**).

### scATAC-seq analysis

10x Illumina data were processed using Cell Ranger ATAC (v.3.1.0) with default parameters and reads were aligned to the human reference sequence GRCh38. All samples were loaded via the Signac package (v1.9.0), then QC filtered based on fragment count vs. TSS enrichment plots to remove low-quality cells and outliers (see **Supp. Fig. 8**). Cells with fewer than 500 fragments, fewer than 30% of reads in cellranger-identified peaks, a nucleosome signal greater than 5, or a percent of blacklisted fragments greater than 5% were removed (the only exception was sample PT-19, which was significantly lower-quality, where the reads in peaks threshold was set to 15%). Doublets were identified using Amulet (v1.1) and removed. Gene scores were calculated using the ArchR (v1.0.2, used in a separate environment with R 4.0.5) package^48^ and imported into Signac^74^. Cell type labels were assigned in the ATAC data using bridge integration^46^, where labels from an annotated scRNA-seq bone marrow reference were assigned using a 10x Multiome bone marrow data set as a “bridge”. Cluster identities were defined first by majority bridge integration cell type label, then verified via chromVAR (v1.20.0) accessibility score for known marker TFs and surface ADT expression. UBA1 genotyping data were processed using GoT-ChA pipeline as described^23^, using KDE (Kernel Density Estimation) applied to log-read distributions to define the boundaries between background noise and genotyping signal for wild-type and mutant reads followed by clustering on z-score read counts.

### Differential motif accessibility analysis

The HSC cluster was isolated and then peak calling was performed again using macs2 (v.2.2.7.1). chromVAR (v1.20.0) was then used to annotate motif enrichment per cell for all motifs in the JASPAR2020 database using the new peak set. TFs that were expressed in fewer than 5 cells in the HSC cluster of our RNA data set were excluded. Samples that had fewer than 5 WT or fewer than 5 MUT cells were then excluded from analysis, leaving 5 samples remaining (from patients PT08, PT10, PT16, PT17, PT18). Linear mixture models were used as previously described^23^ to determine significantly differentially accessible motifs while controlling for patient-specific effects.

### Differential gene score analysis

The HSC cluster was isolated, and samples that had fewer than 5 WT or fewer than 5 MUT cells were then excluded from analysis, leaving 5 samples remaining (from patients PT8, PT10, PT16, PT17, PT18). Because of the sparsity of estimating individual gene expression data per cell in ATAC data sets, we applied extra filtering to retain only high-quality cells, filtering for number of ATAC fragments > 1,500 and percent reads in peaks > 50%. Genes that were expressed in fewer than 20% of the cells in the HSC cluster of our corresponding RNA data were excluded, as were cells with gene expression scores in fewer than 20% of our ATAC HSCs. Linear mixture models were used as previously described^23^ to determine significantly differentially accessible motifs while controlling for patient-specific effects. Gene set enrichment analysis was performed as described in the RNA differential expression analysis (using the same set of 95 gene modules, ranked by effect size using the fgsea package, Benjamini-Hochberg correction used).

### ADT processing

All ADT libraries were processed using kallisto/bustools. We used the dsb package^75^ (v1.0.3) as an alternative form of normalization for the ADT protein expression values. Normalized values were applied for selection filtering of ADT markers for which the true signal was above the background noise levels, within the captured cell-contained droplets. dsb discriminates between background noise by differentiating between empty droplets (containing ambient mRNA and antibody, but no cell) and true cell-containing droplets. The background matrix was defined from the comparison of the raw feature barcode matrices from the 10x sequencing output versus the processed filtered feature barcode matrix results generated from running Cell Ranger (see above). The final output filters out empty droplets and retains only true cell-containing droplets based on the 10x cell calling algorithm. As such, the matrix of background noise is generated by subtracting out the positive cell containing droplets found in the filtered matrices from the negative empty droplets in the raw matrices. Normalization was performed using the DSBNormalizeProtein function. The dsb normalized values were defined as the number of standard deviations above the background noise. For the RNA ADT data, differential ADT expression testing was performed using linear mixture models as previously described on the dsb-corrected values.

### Phospho-seq analysis

Antibodies used in the TSA phospho-seq panel were quantified via alevin from the package salmon (1.10.1) using the same parameters as the original publication: --read-geometry 2[11-25] --bc-geometry 1[1-16] --umi-geometry 2[1-10] (see **Supp. Table 18** for antibody list). Intracellular antibody counts were then merged with the scATAC-seq data using shared cell barcodes, and normalized using CLR normalization using the same workflow as is recommended by Seurat for surface protein data. To generate rank plots, the HSC, EMP, LMPP, and MkP clusters were isolated and the difference of the mean expression values in MUT and WT cells was calculated for each protein, which was then used for ranking.

### Control sample analysis

Control samples from healthy donors (HD1_CD34, HD2_CD34, HD3_CD34, HD4_CD34, HD1_BM, HD2_BM, HD3_BM, HD4_BM) were downloaded from the Gene Expression Omnibus (GEO) database accession GSE196052. Raw fastq files were aligned to the human reference sequence GRCh38 using Cell Ranger (v.3.1.0) with default parameters. Ambient mRNA was removed from each sample using SoupX software (v1.6.2) with default parameters. Quality control was performed to remove low-quality cells exhibiting a percentage of mitochondrial reads above 10%. Potential doublets exhibiting a number of unique expressed genes above 3,000 were removed as well as empty droplets of less than 500 unique expressed genes. The gene per cell matrix of each sample was log-normalized using the Seurat LogNormalize method and scaled regressing-out the percentage of mitochondrial reads, the S phase score and the G2/M phase score calculated for each cell using the CellCycleScoring Seurat function. The scaled matrix was then used for Principal Component Reduction (PCA) using the RunPCA function from the Seurat package with seed.use parameter set to 1 for reproducibility.

### Control sample integration and transfer label

To mitigate the potential heterogeneity between samples introduced by different processing methods, we used the Seurat workflow to project healthy donor cells onto the VEXAS mRNA UMAP embedding. We first used the FindTransferAnchors function on log-normalized mRNA data using 50 principal components to project the PCA structure of the VEXAS cells onto the healthy donor samples and identify mutual nearest neighbors (anchors) between the datasets. Then, we used the MapQuery to both transfer cell type assignment from the reference VEXAS cells to the healthy donor cells based on a weighted vote classifier, and project the healthy donor data onto the same original UMAP embedding. The query.dims and ref.dims parameters of the MapQuery function were respectively set to 1:30 and 1:50.

Finally, we merged the data from both VEXAS and the healthy donors into a final Seurat object for downstream analysis. This unified object ensured that all cells were mapped onto the same UMAP embedding, with healthy donor cells assigned to the distinct cell types.

### Control sample identification of differentially expressed genes

We performed transcriptome-based differential expression tests for the different cell types between VEXAS and healthy donor cells. We first removed all mitochondrial and ribosomal genes, and then reran log-normalization using only the remaining genes. To limit potential sample-specific cofounders, we randomly selected 200 cells from each sample within each specific cell type. We considered all genes detected in at least 10% of the VEXAS cells with a log2 fold change larger than 0.5 or smaller than −0.5 and a Bonferroni-adjusted *P* < 0.05 as differentially expressed.

### Control sample gene module score enrichment analysis

HALLMARK_APOPTOSIS, HALLMARK_INFLAMMATORY_RESPONSE, GOBP_COMMON_MYELOID_PROGENITOR_ CELL_PROLIFERATION, REACTOME_UNFOLDED_PROTEIN_RESPO NSE_UPR gene sets were imported using the msigdbr R package, providing Molecular Signatures Database (MSigDB) gene sets typically used with the Gene Set Enrichment Analysis (GSEA) software.

We calculated module scores of these four gene sets for each single cell using the AddModuleScore function from Seurat. Briefly, the function infers the gene set activity by calculating the average expression levels of each gene set on a single-cell level, subtracted by the aggregated expression of control feature sets. All analyzed features are binned based on averaged expression, and the control features are randomly selected from each bin. Default parameters were used and the seed set to 1234 for reproducibility.

To plot the myeloid differentiation gene set enrichment in VEXAS vs. in control HSCs, the GOBP_COMMON_MYELOID_PROGENITOR_ CELL_PROLIFERATION pathway was used to assess myeloid markers activity in each cell (as detailed above) and kernel density estimate for both VEXAS and control cells were calculated and visualized using the ggplot2 R package and the geom_density function using a multiplicate bandwidth adjustment of 1.5.

### Pseudotime analysis

Pseudotime cell ordering was performed using the Monocle3 (version v1.3.1) workflow in a semi-supervised manner. First, cells were divided into two lineages: myeloid (HSC, LMPP, GMP, CD14 Mono, MkP, and EMP) and lymphoid (HSC, LMPP, NK, CLP, B, CD4 T, and CD8 T). The cluster_cells function was then applied with default parameters to cluster the cells within each lineage. Next, the learn_graph function was used to learn the principal graph connecting the cell clusters, with the use_partition parameter set to false. Finally, the order_cells function was used to assign a pseudotime to each cell based on their projection onto the learned principal graph and their distance from the root HSC cells at the top of the trajectory graph.

## Supplementary figures

**Supplementary Figure 1.**
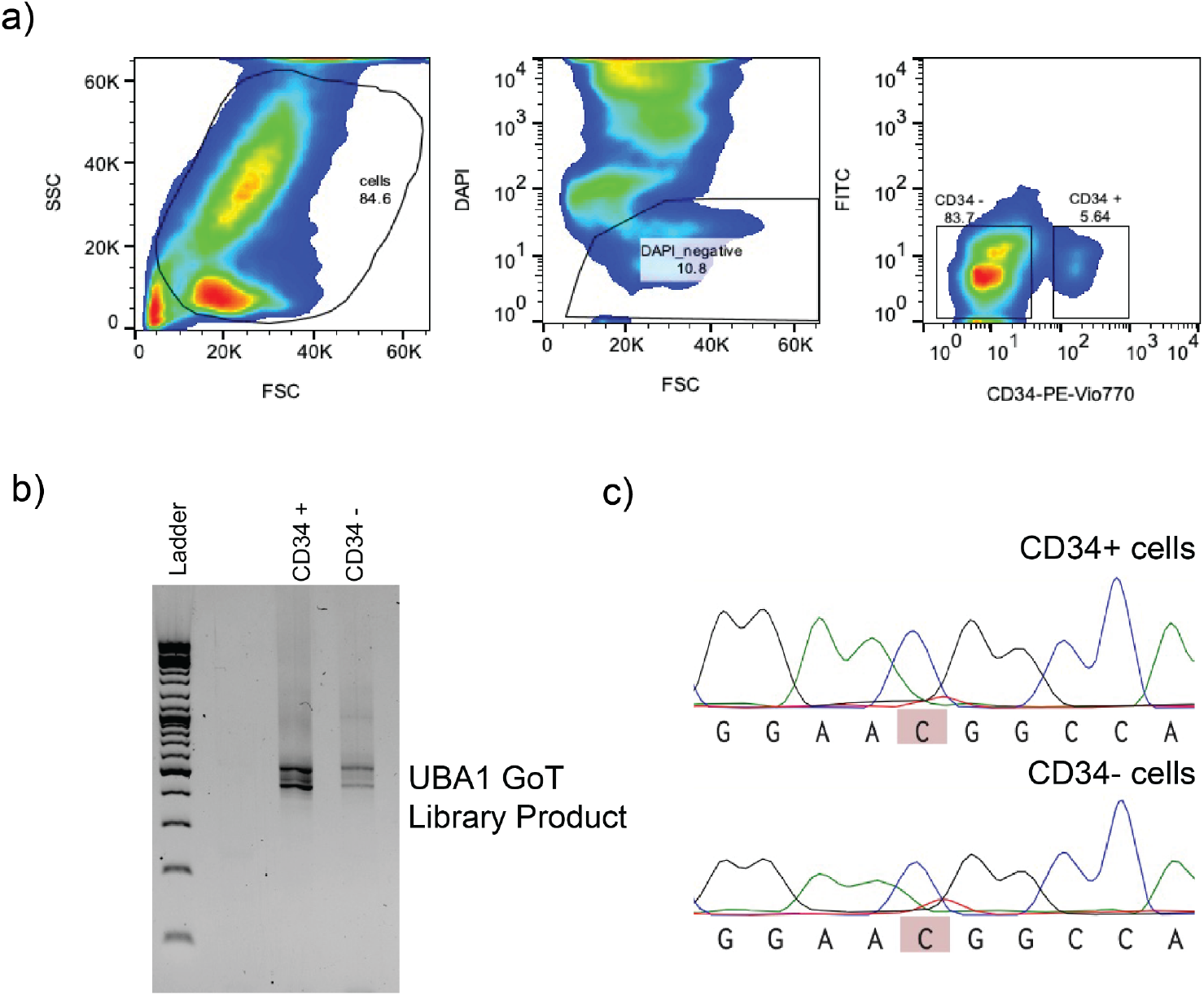
Representative images of **(a)** CD34+ cell sorting strategy. **(b)**UBA1 GoT library final product on 2% agarose gel performed on both CD34+ and CD34-compartment sorted cells. **(c)** Confirmation of UBA1 p.M41T (c.122 T>C) genotype from GoT library by Sanger sequencing. The altered base is highlighted showing >90% VAF.

**Supplementary Figure 2.**
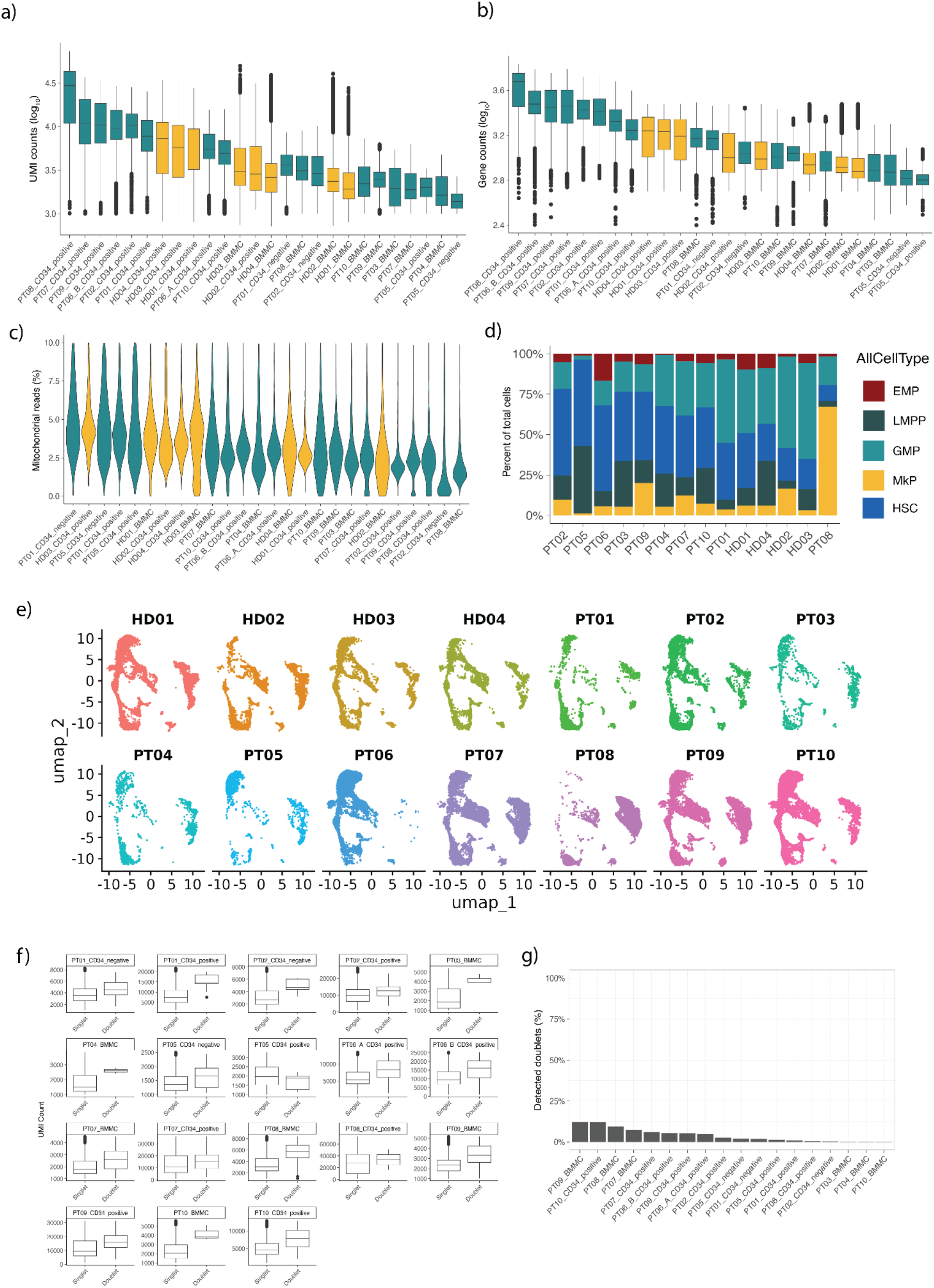
scRNA-seq data QC and doublet removal. **(a-c)** Distribution of UMI count per cell (a), gene count per cell (b), and percent mitochondrial reads (c) post-QC filtering for all RNA samples analyzed, VEXAS (green) and healthy controls (yellow). Error bars represent the range, boxes represent the interquartile range and lines represent the median. **(d)** Percent of cells mapping to each progenitor sub-type post cell type assignment. **(e)** UMAPs plotted for each individual sample, showing consistent distribution of cells after reciprocal PCA integration (Methods). **(f-g)** Doublet removal results showing the UMI count per cell between labeled doublets and singlets (f) and percent of cells in each sample labeled doublets (g). Doublet removal was performed using DoubletFinder (Methods).

**Supplementary Figure 3.**
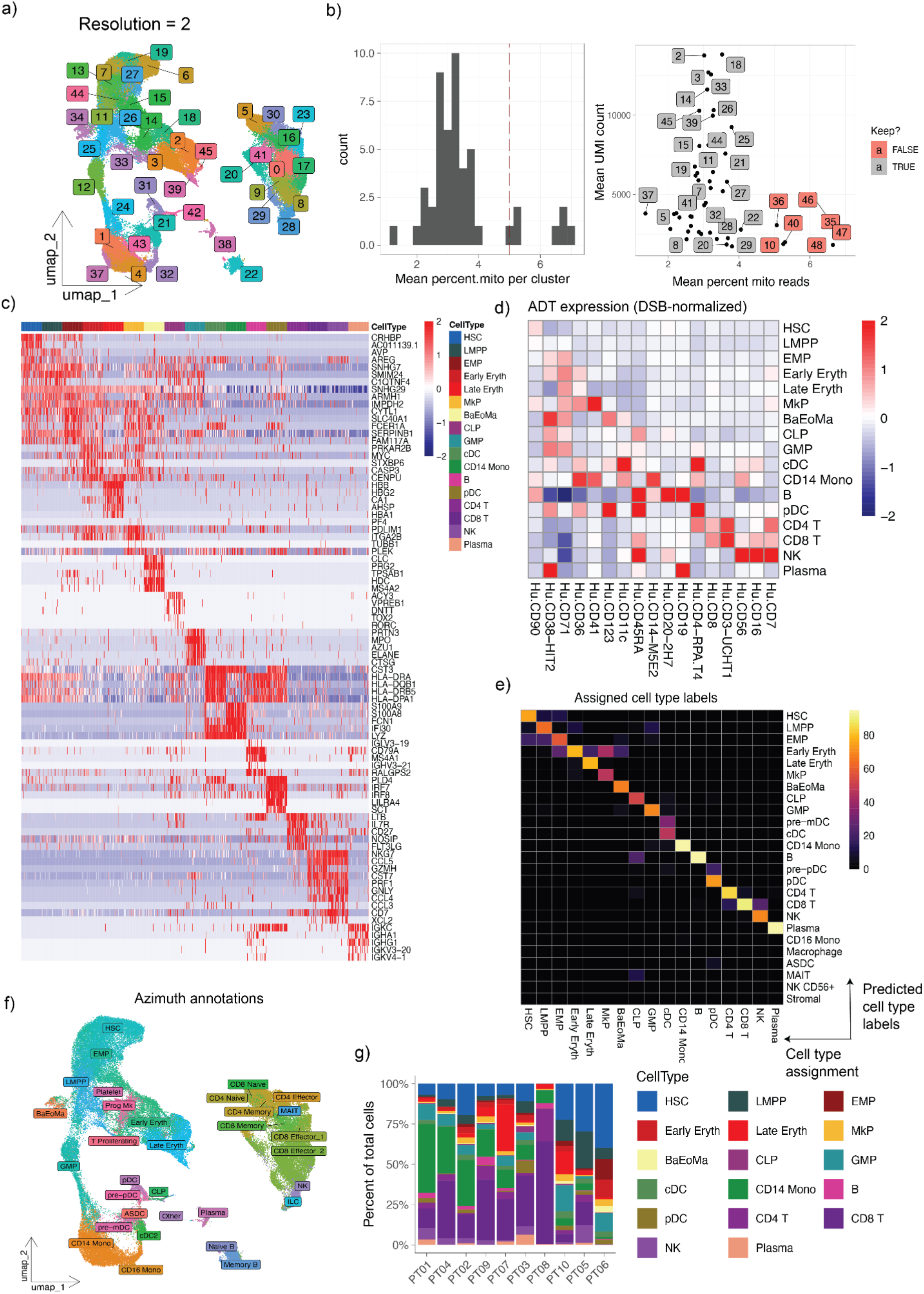
scRNA-seq clustering and cell type annotation. **(a)** Results from cluster identification using the FindNeighbors and FindClusters functions in Seurat. **(b)** Identification of outlier clusters in terms of mean percent mitochondrial reads (left) as well as mean UMI counts (right), representing low-quality clusters that were excluded from subsequent analysis. Red dashed line in left panel indicates threshold used to remove outliers. **(c)** Heatmap showing the top 5 marker genes of each annotated cell type. **(d)** Heatmap showing surface protein expression of known cell type markers for each annotated cell type. **(e)** Heatmap showing the alignment between Azimuth cell type labels and annotated cell types. **(f)** UMAP annotated with Azimuth cell type labels. **(g)** Bar plot showing the breakdown of cell types identified from each donor.

**Supplementary Figure 4.**
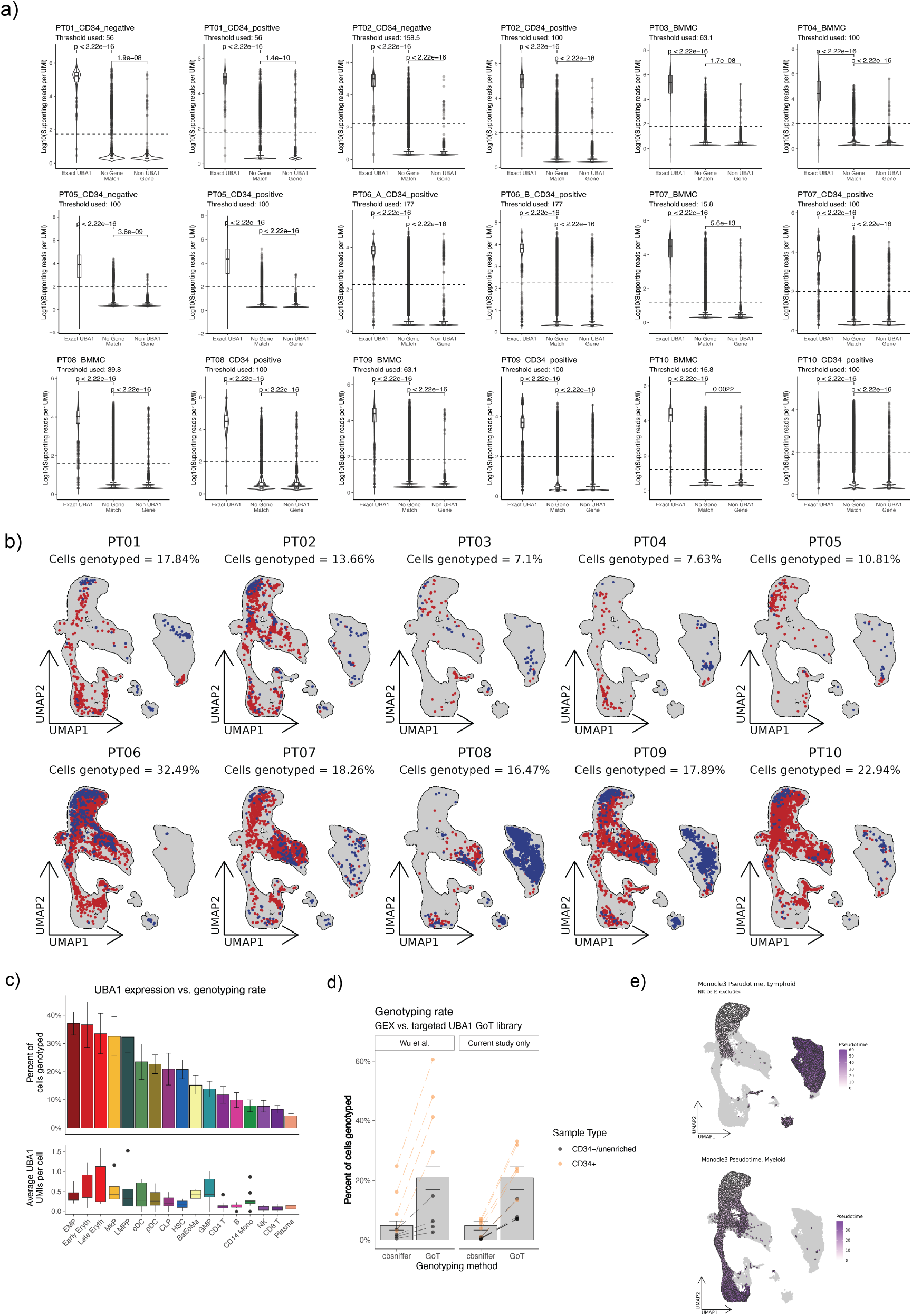
GoT genotyping data analysis. **(a)** Distributions of reads per UMI found in the genotyping library when separated by UMIs that matched the target gene (*UBA1*) in the gene expression library, UMIs that matched a non-target gene that are likely products of PCR recombination, and UMIs with no match, across patient samples. Cutoffs were selected based on the local minimum of the bimodal no-match distribution. Two-sided Wilcoxon rank sum test. **(b)** UMAPs annotated with genotyping information (blue = WT; red = mutant; grey = NA), for each patient. **(c)** Comparison of genotyping rate (top) with *UBA1* expression (bottom), as a function of cell type. Bar plot: error bars represent standard error of the mean. Box plots: Error bars represent the range, boxes represent the interquartile range and lines represent the median. Asterisks represent outliers. **(d)** Comparison of genotyping rates obtained from using GoT vs. only using the existing 10x 5’ biased data (cbsniffer). Error bars represent standard error of the mean. **(e)** Pseudotime analysis of lymphoid and myeloid trajectories.

**Supplementary Figure 5.**
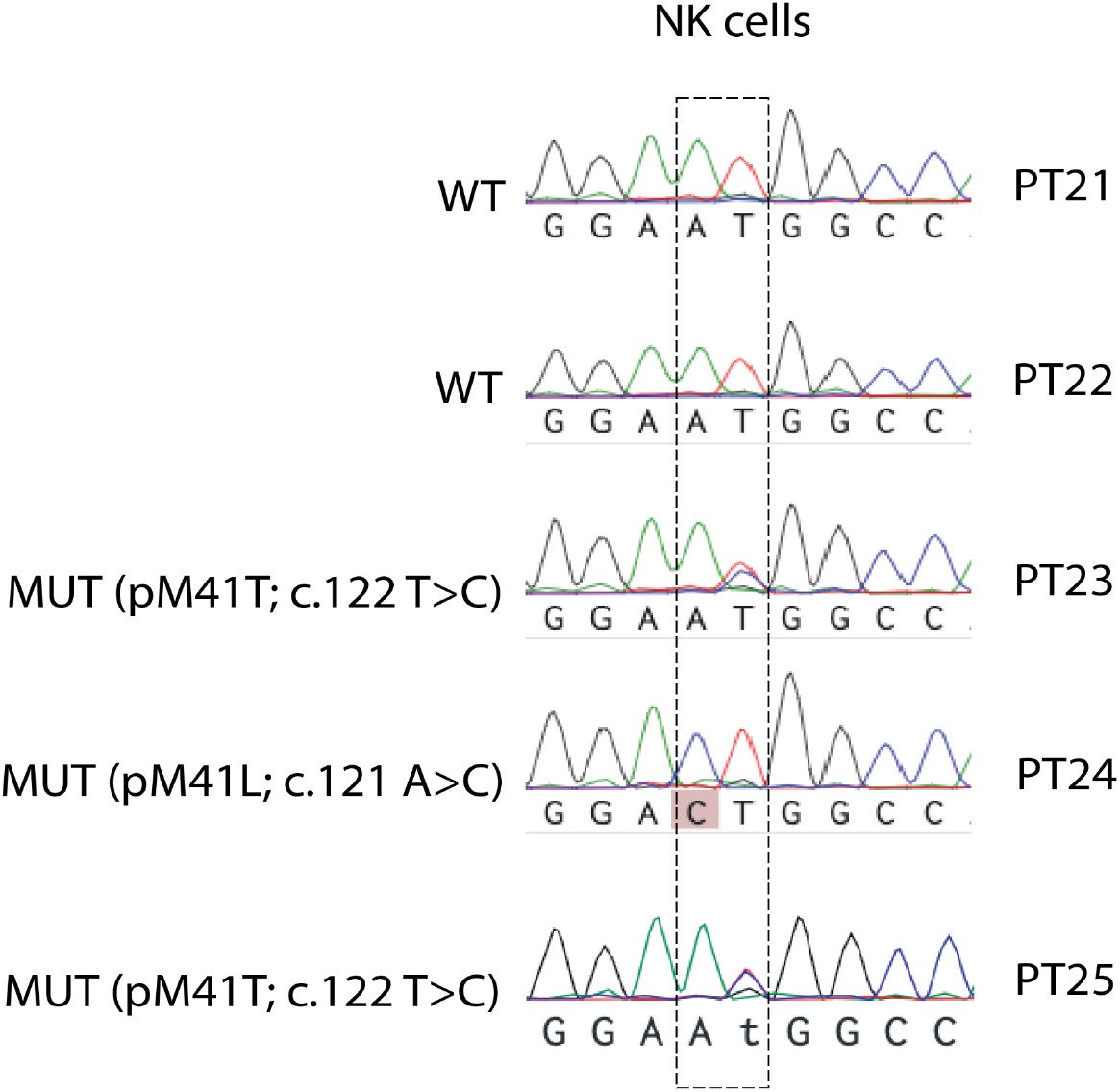
Validation of *UBA1* mutation in NK cells from VEXAS patients. PT21, PT22 show absence of *UBA1* mutation in NK cells. PT23-PT25 shows the presence of *UBA1* mutation in NK cells.

**Supplementary Figure 6.**
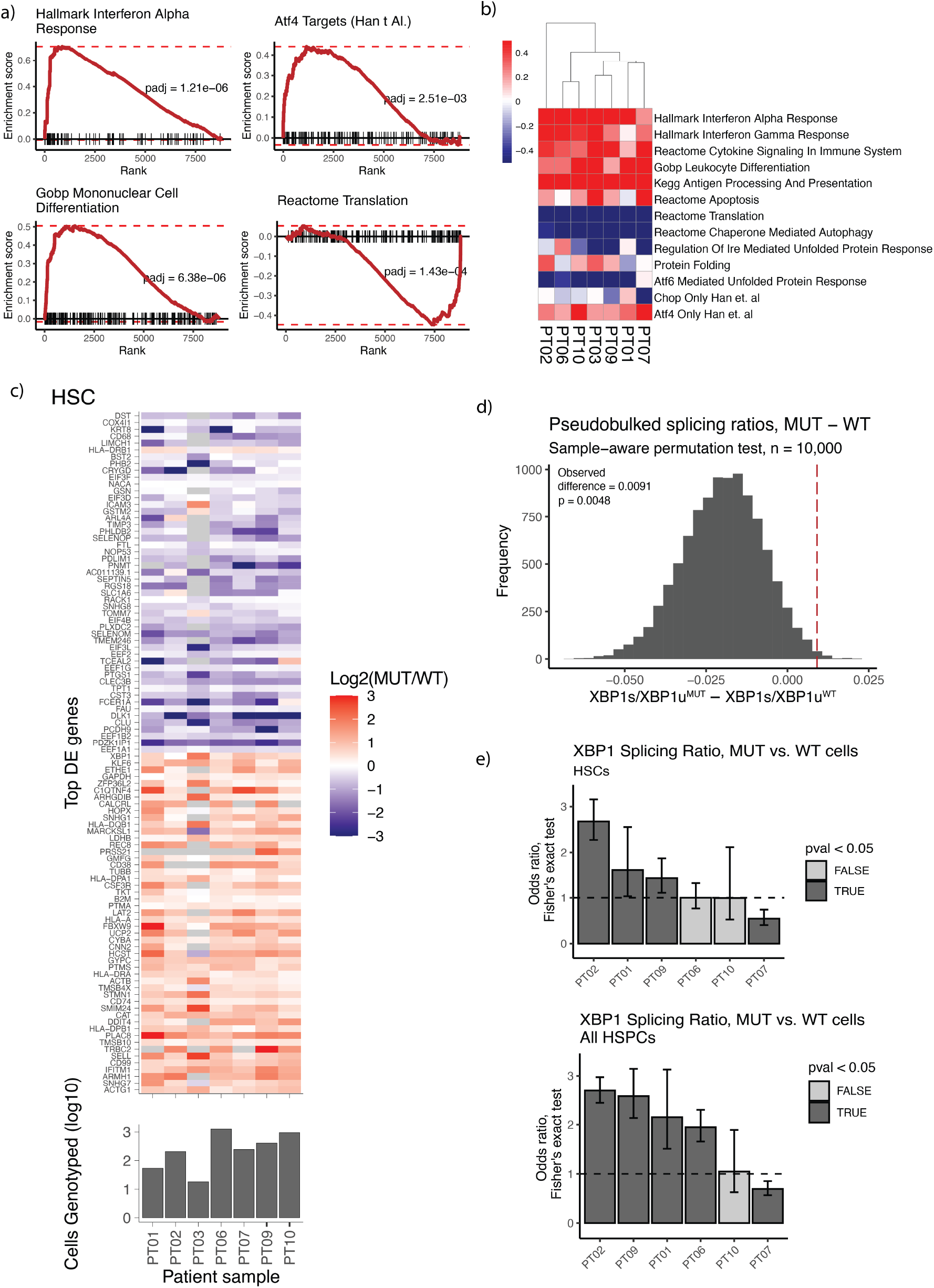
Comparison of *UBA1*-mutant vs. WT cells in scRNA-seq data. **(a)** Ranked gene set enrichment analysis (GSEA) of pathways of interest from the differential expression analysis between MUT and WT HSCs (Supplementary Table 7). **(b)** Heatmap of GSEA enrichment score in mutant HSCs across seven patients. Color scale indicates the mean enrichment score difference between *UBA1*-mutated and wild-type HSCs. **(c)** Heatmap of Top DE genes in mutant HSCs across seven patients. Color scale indicates the log2-fold change between the average expression in *UBA1*-mutated and wild-type HSCs. **(d)** Distribution of sample-aware permutations (mutant and wild-type labels permuted only within sample) comparing XBP1 spliced / XBP1 unspliced ratios in MUT and WT HSCs (Methods). Dashed red line is the observed value. **(e)** Comparison of spliced (XBP1s) to unspliced (XBP1u) *XBP1* detected in *UBA1*-mutant versus wild-type HSCs, pseudobulked by genotype and patient. Only patients with sufficient number of genotyped HSCs (upper panel) and HSPCs (lower panel) (>= 25 MUT, >= 25 WT) are shown. P-value determined by Fisher’s exact test. Bars represent the odds ratio, and error bars represent the 95% confidence interval. Dotted line indicates odds ratio of 1.

**Supplementary Figure 7.**
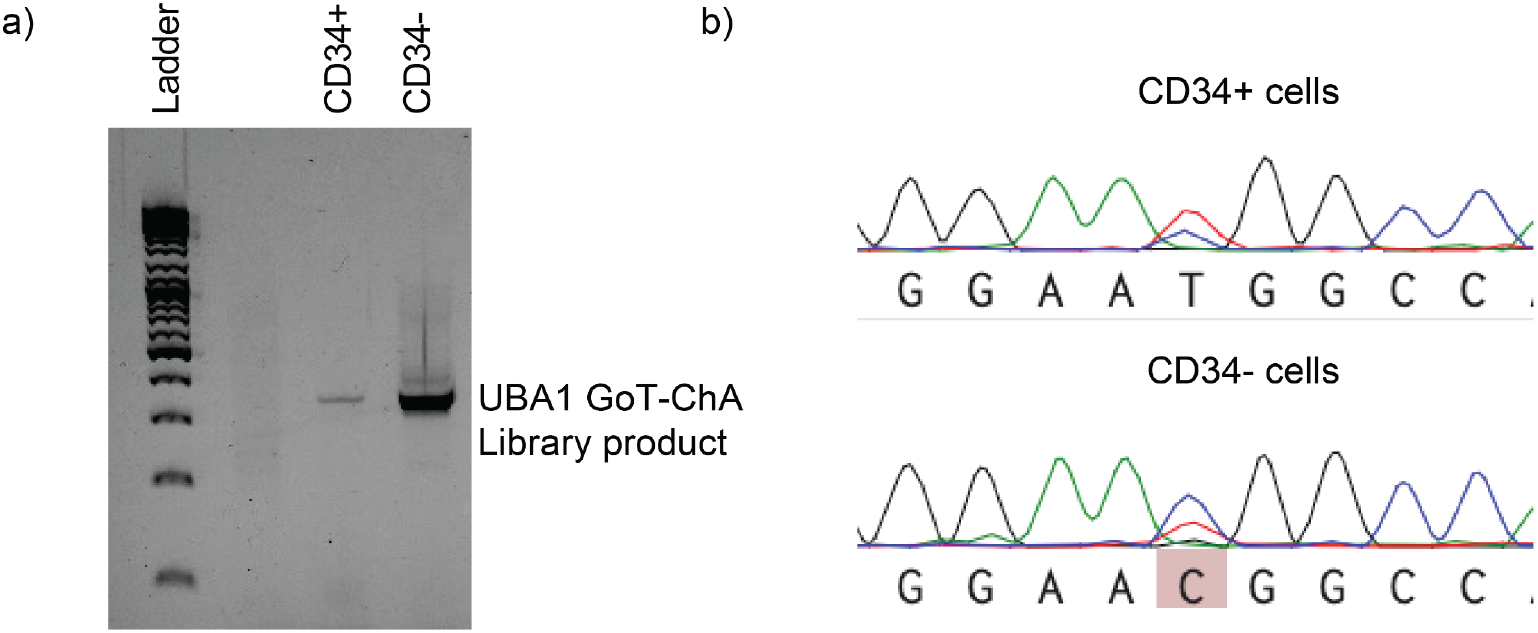
Representative images of **(a)** UBA1 GoT-ChA library final product on 2% agarose gel performed on both CD34+ and CD34-compartment sorted cells. **(b)** Confirmation of UBA1 p.M41T (c.122 T>C) genotype from GoT-ChA library by Sanger sequencing. The mutation is highlighted.

**Supplementary Figure 8.**
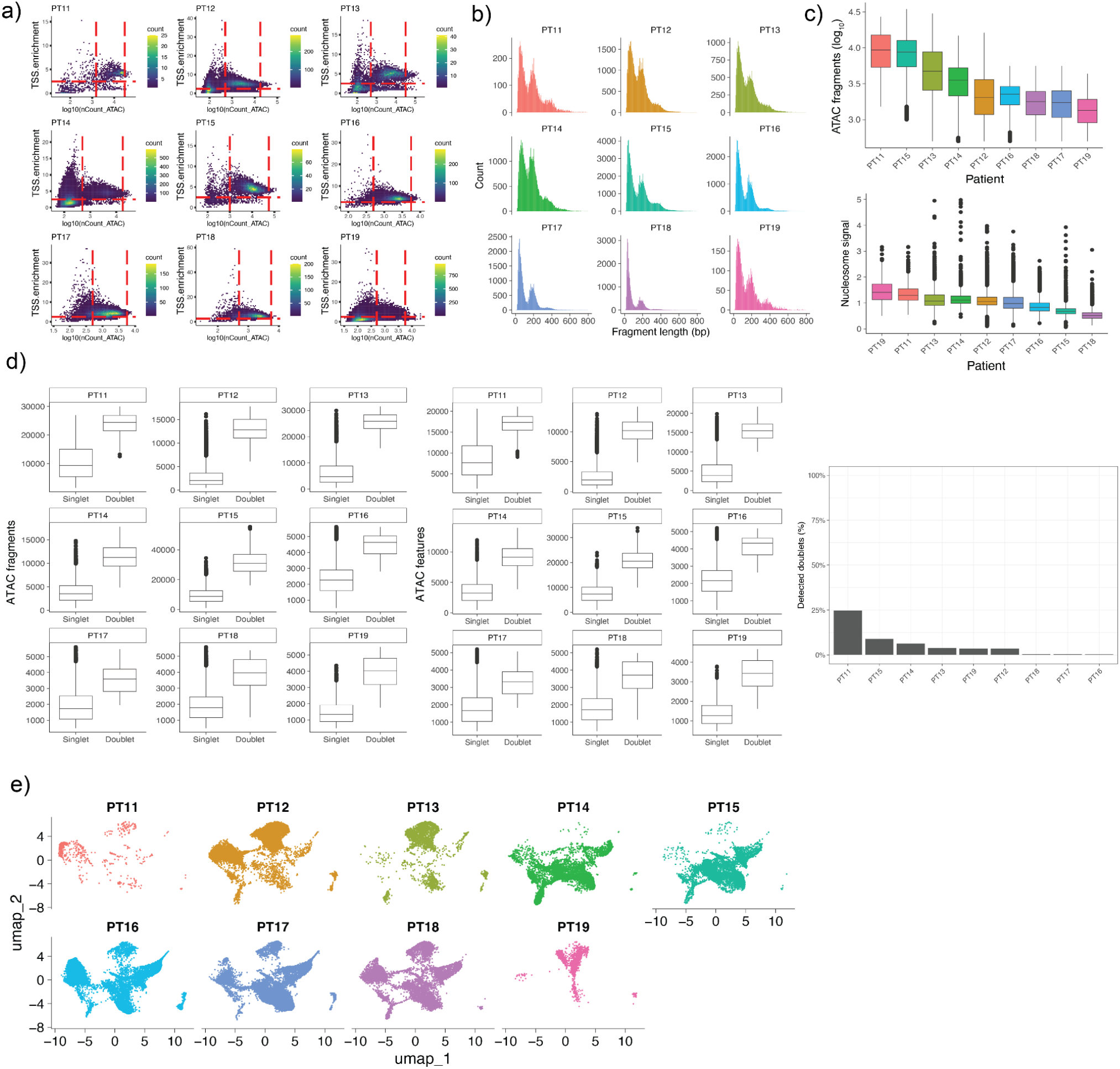
QC and doublet removal in scATAC-seq data. **(a)** Scatter plots of read count per cell vs. transcription start site (TSS) enrichment in each scATAC-seq sample. Dashed red lines indicate thresholds used for QC filtering. **(b)** Histogram plots showing fragment length (bp) enriched in each sample. **(c)** Box plots of number of reads (top) and of nucleosome signal (bottom) in each scATAC-seq sample. Error bars represent the range, boxes represent the interquartile range and lines represent the median. **(d)** Doublet removal results showing the read count per cell between labeled doublets and singlets and percent of cells in each sample labeled doublets. Doublet removal was performed using Amulet (Methods). Error bars represent the range, boxes represent the interquartile range and lines represent the median. **(e)** Accessibility-based UMAP plots split by patient showing consistent distribution of cells after reciprocal LSI integration (Methods).

**Supplementary Figure 9.**
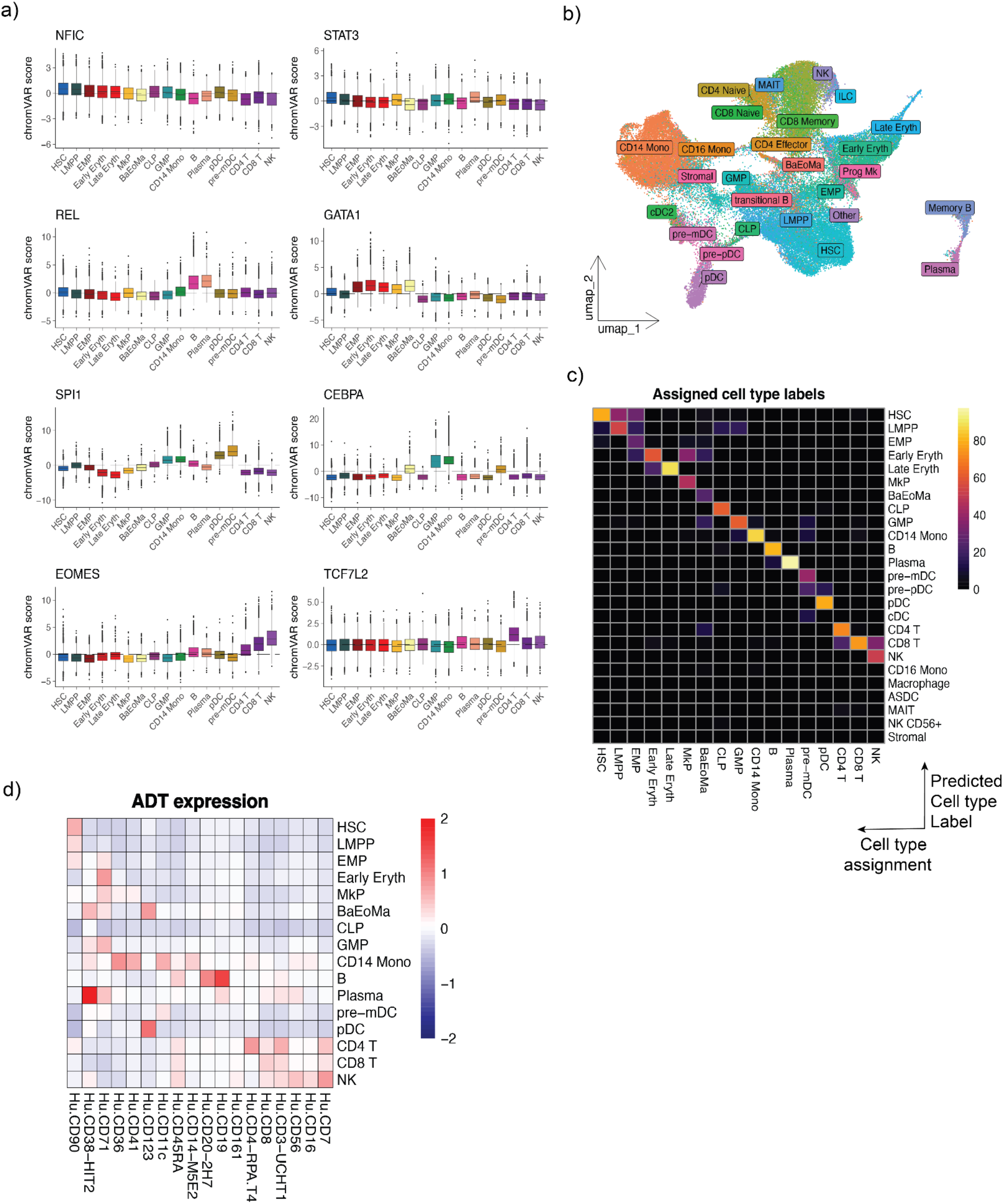
Cell type annotation for scATAC-seq data. **(a)** Enrichment of marker TF motifs as a function of annotated cell type. Error bars represent the range, boxes represent the interquartile range and lines represent the median. **(b)** UMAP annotated with cell type labels, assigned from an annotated bone marrow scRNA-seq data set via bridge integration (Methods). **(c)** Heatmap showing the alignment between bridge integration cell type labels and annotated cell types. **(d)** Heatmap with surface protein expression of canonical cell type markers per annotated cell type.

**Supplementary Figure 10.**
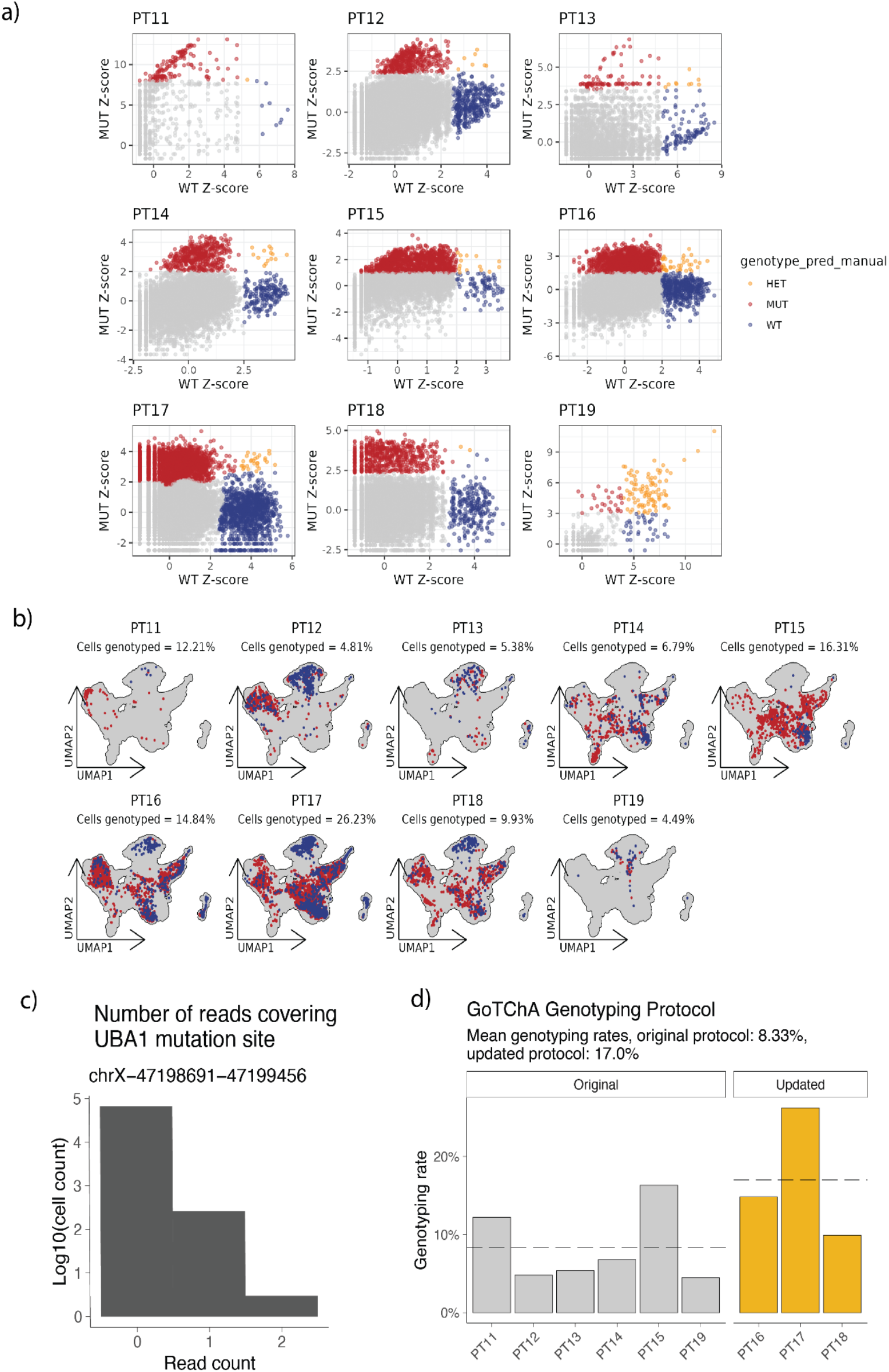
Genotyping via GoT-ChA. **(a)** Transformed distribution of MUT and WT reads per cell (Methods). Color indicates resulting genotype assignment (Red = MUT, blue = WT, yellow = HET excluded as doublets). Cells with a genotype assignment of HET were assumed to be doublets and excluded from further analysis. **(b)** UMAPs annotated with genotyping assignments (blue = WT; red = mutant; grey = NA) for each patient. **(c)** Distribution of read counts per cell in the original ATAC-seq library overlapping the *UBA1* mutation site. Most cells had no coverage, and the majority of those with coverage had a single read. **(d)** Bar plot of overall genotyping rate per sample. Samples from patients 16, 17, and 18 were processed using an updated GoT-ChA protocol with increased PCR annealing time that improved genotyping rate. Horizontal dashed line: mean genotyping fraction among samples treated with original protocol and updated protocol.

**Supplementary Figure 11.**
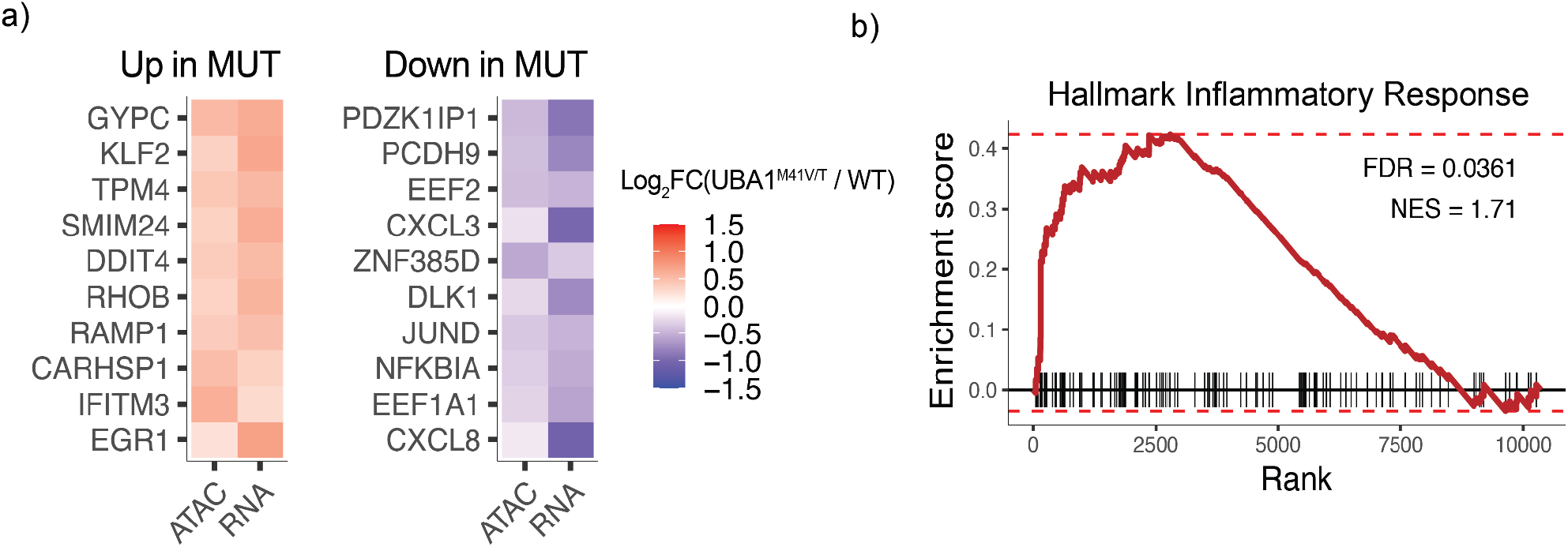
**(a)** Heat map showing the log2FC of genes identified as significant in both the RNA differential expression analysis and the ATAC gene score differential accessibility analysis. Genes included in both analyses were ranked by multiplying log2FC^RNA^*log2FC^ATAC^*sign(log2FC^RNA^), and the top 10 and bottom 10 genes are shown. **(b)** Gene set enrichment analysis of genes with high differential accessibility scores showing enrichment of Hallmark Inflammatory Response pathway. Preranked gene set enrichment analysis was performed using the fgsea package.

**Supplementary Figure 12.**
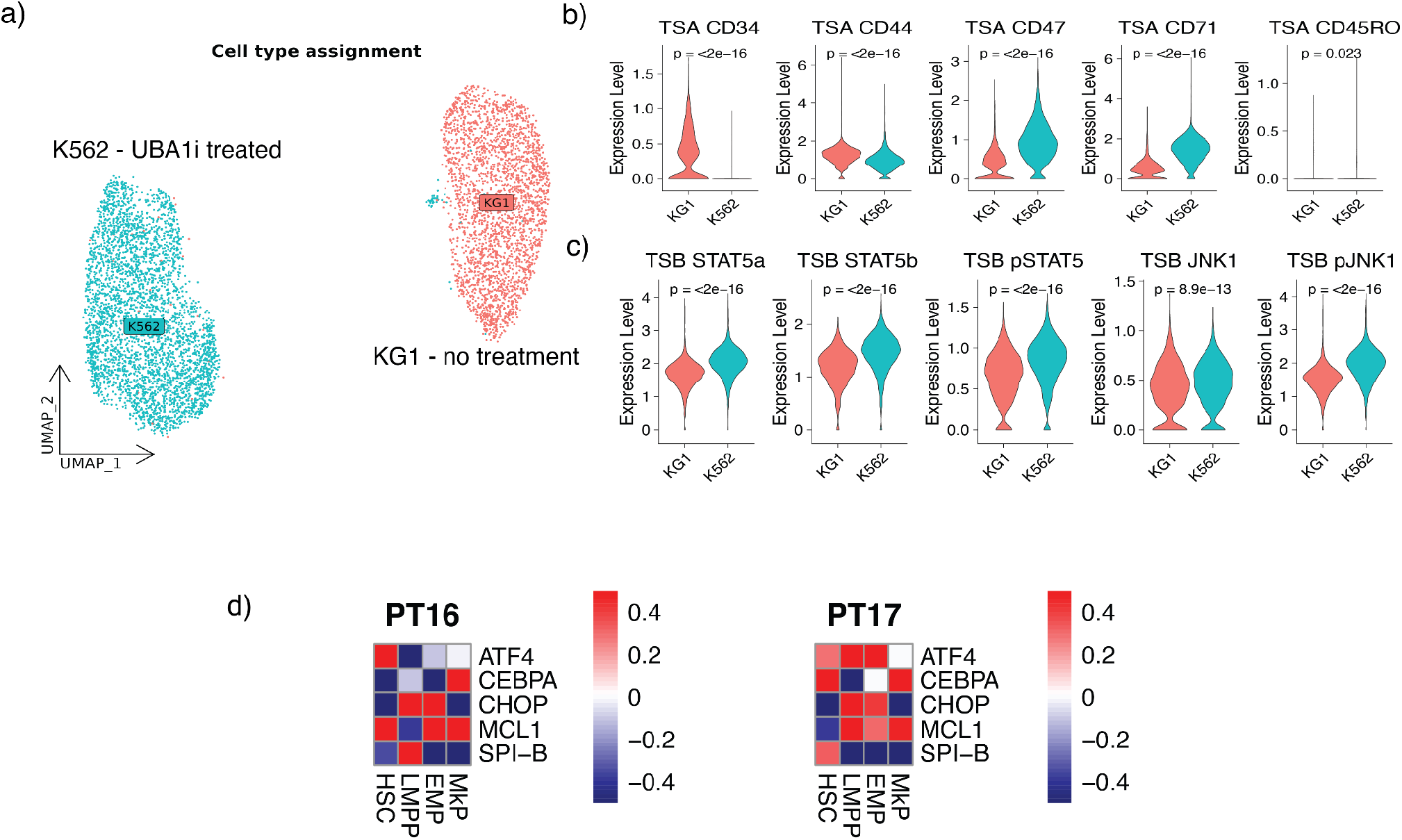
Phospho-seq cell line mixing experiment. **(a)** UMAPs of scATAC-seq data from a cell line mixing experiment to test the custom Phospho-seq intracellular antibody panel. The two cell lines, K562 and KG1, are clearly separated by their epigenetic profiles. The K562 cell line was also exposed to the UBA1 inhibitor TAK-243 to mimic the *UBA1* mutation. **(b)** Surface protein data showing that the K562 and KG1 clusters show the expected cell surface markers (CD34, CD44 up in KG1, while CD47, CD71 are up in K562). Two-sided Wilcoxon rank sum test. **(c)** Intracellular protein data showing that the K562 cells have higher quantities of STAT5 (expected due to the documented BCR-ABL fusion in this cell line) as well as of JNK1 (expected due to the UBA1 inhibition). Two-sided Wilcoxon rank sum test. **(d)** Heatmaps of average expression of targeted proteins in two different patients (PT16 and PT17) showing different HSPC progenitors in MUT vs. WT (scaled by cell type).

**Supplementary Figure 13.**
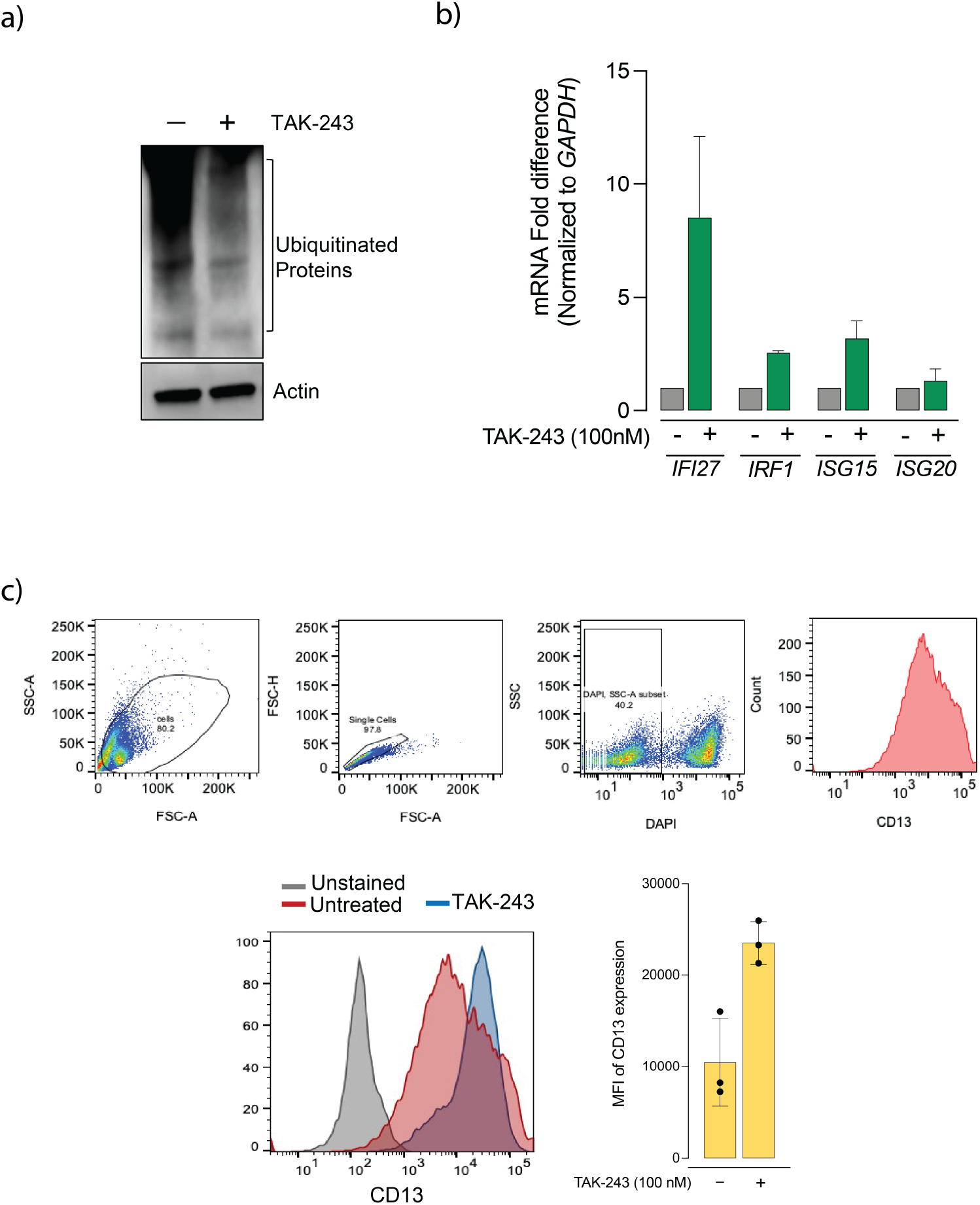
TAK-243 treatment induces VEXAS like phenotype in CD34+ cells ex vivo. **(a)** Western blot showing reduced ubiquitination of proteins when CD34+ cells were treated with TAK-243 for 48 hours (representative image). **(b)** Q-PCR results show increased expression of inflammatory genes upon treatment with TAK-243 for 48 hours. Error bars represent standard error. **(c)** Gating strategy for CD13 expression (myeloid marker) on CD34+ cells subjected to myeloid differentiation upon TAK-243 treatment showing increased expression of CD13 marker (n = 3 donors). Error bars represent standard error. MFI, mean fluorescence intensity.

**Supplementary Figure 14.**
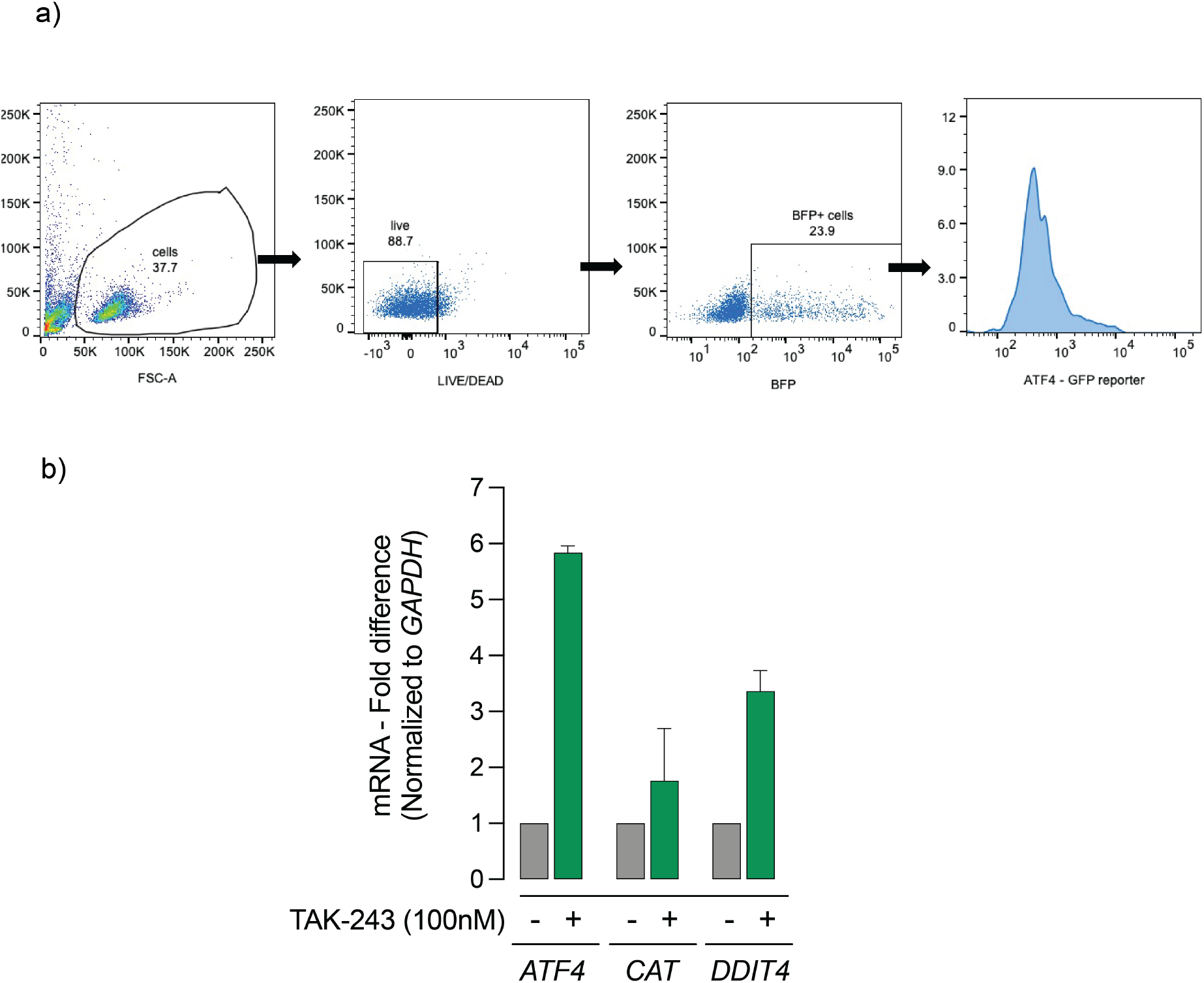
**(a)** Gating strategy used for ATF4 reporter assay analysis. **(b)** Expression levels of *ATF4, CAT* and *DDIT4* genes normalized to *GAPDH* expression measured by quantitative RT-PCR in TAK-243-treated and untreated CD34^+^ cells from Donor 1 performed in technical duplicates. Error bars represent standard error.

**Supplementary Figure 15.**
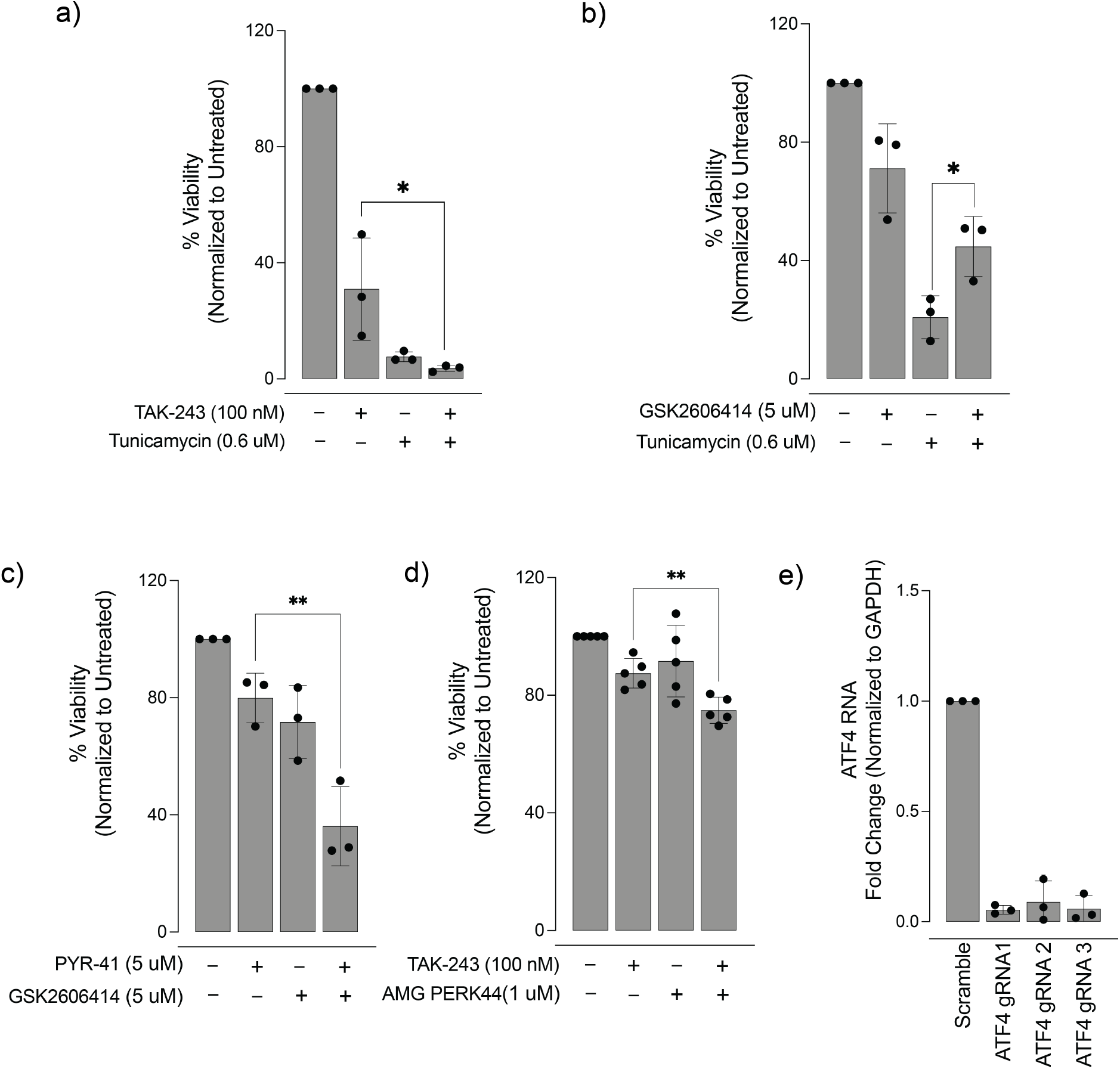
**(a-d)** Apoptosis assay showing, **(a)** TAK-243 along with tunicamycin reduces the viability of CD34+ cells (96 hours). **(b)** PERK inhibitor (GSK2606414) rescues tunicamycin-induced apoptosis in CD34+ cells. **(c)** UBA1 inhibitor (PYR-41) along with PERK inhibitor (GSK2606414) reduces the viability of CD34+ cells. **(d)** UBA1 inhibitor (TAK-243) along with another PERK inhibitor (AMG PERK44) reduces the viability of CD34+ cells. **(e)** Q-PCR analysis showing efficient knockout of *ATF4* in 3 different donors compared to scramble. Error bars represent standard error. ** indicates significance, two-sided Student t-test.

**Supplementary Figure 16.**
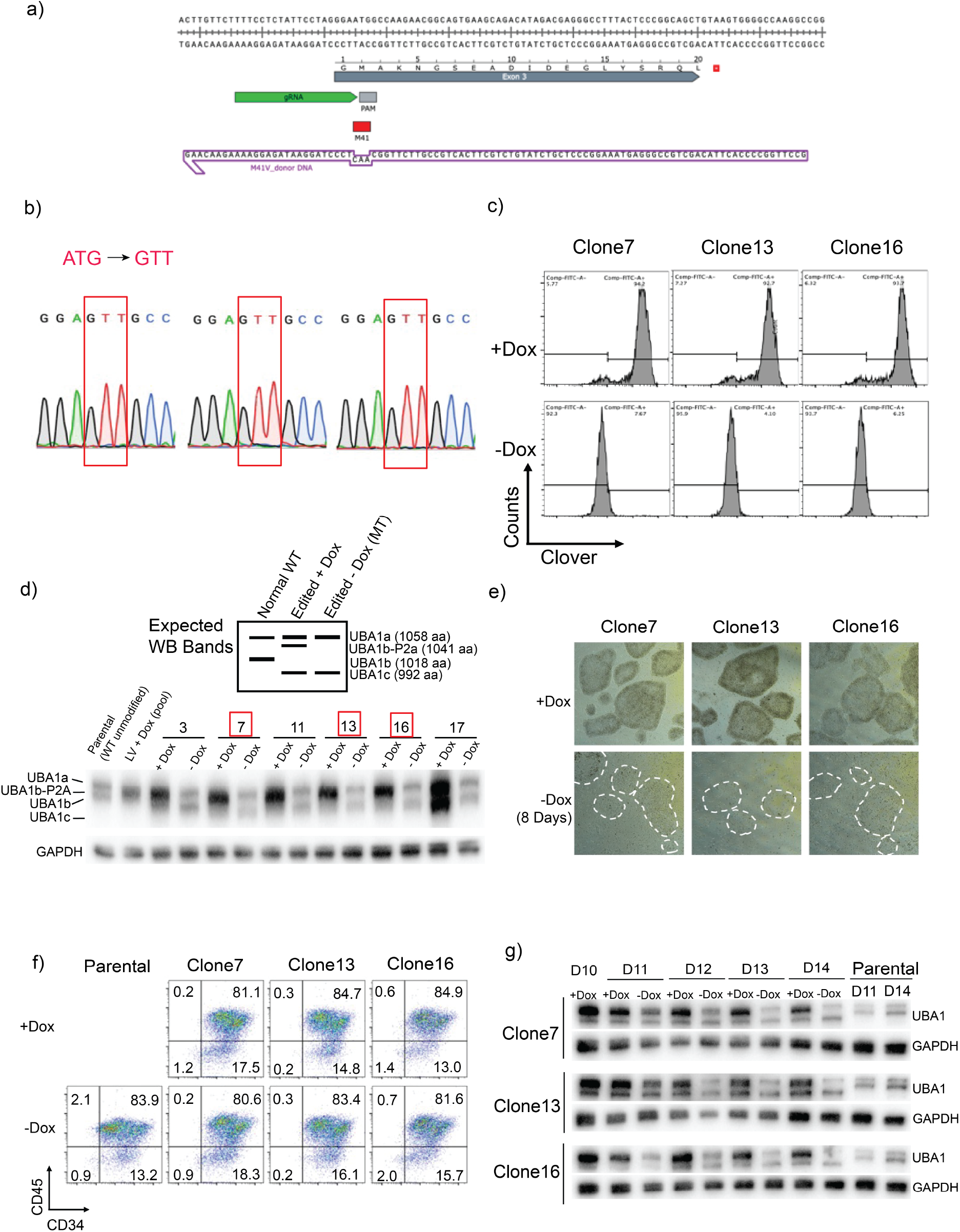
**(a)** Gene editing strategy. Sequence of the *UBA1* locus with the positions of the gRNA and M41V mutant donor DNA indicated. **(b)** Sanger sequencing of the edited region in 3 selected clones. Red boxes indicate the M41 ATG edited to GTT (V). **(c)** Flow cytometry of iPSCs from the 3 selected clones in the presence of doxycycline (upper panels) or after withdrawal of doxycycline (lower panels), showing expression of Clover, co-expressed with UBA1b. **(d)** Western blot analysis of UBA1 isoforms in edited iPSC clones (indicated by numbers) in the presence or absence of Dox. The expected band pattern in wild-type (WT), edited and mutant cells is shown in the upper panel. **(e)** iPSC colonies from 3 edited clones grow normally in the presence of Dox, but not without, indicating dependence on expression of the UBA1b isoform. **(f)** CD34 and CD45 expression in HSPCs derived from the parental iPSC line or 3 different VEXAS iPSC clones on day 12 of differentiation. Dox was removed, as indicated, on day 10 of differentiation. **(g)** Western blot analysis of UBA1 isoforms in HSPCs from 3 indicated *UBA1*^*M41V*^-mutant iPSC clones on days 10-14 of differentiation in the presence or absence of Dox, as indicated.

## Notes

### Summary of Updates

Updated the author list and contribution SaG, RMM, JS, EOE, KT, FI, ZW, ShG, NSY, JDB, RS, BT, MH, IR, EPP, PS, OK, DBB and DAL helped interpret results.

